# Targeting Fibroblast-Endothelial Interactions in LAM Pathogenesis: 3D Spheroid and Spatial Transcriptomic Insights for Therapeutic Innovation

**DOI:** 10.1101/2023.06.12.544372

**Authors:** Sinem Koc-Gunel, Emily C. Liu, Lalit K. Gautam, Ben A. Calvert, Shubha Murthy, Noa C. Harriott, Janna C. Nawroth, Beiyun Zhou, Vera P. Krymskaya, Amy L. Ryan

## Abstract

Lymphangioleiomyomatosis (LAM) is a progressive lung disease with limited treatments, largely due to an incomplete understanding of its pathogenesis. Lymphatic endothelial cells (LECs) invade LAM cell clusters, which include HMB-45-positive epithelioid cells and smooth muscle α-actin-expressing LAM-associated fibroblasts (LAMFs). Recent evidence shows that LAMFs resemble cancer-associated fibroblasts, with LAMF-LEC interactions contributing to disease progression. To explore these mechanisms, we used spatial transcriptomics on LAM lung tissues and identified a gene cluster enriched in kinase signaling pathways linked to myofibroblasts and co-expressed with LEC markers. Kinase arrays revealed elevated PDGFR and FGFR in LAMFs. Using a 3D co-culture spheroid model of primary LAMFs and LECs, we observed increased invasion in LAMF-LEC spheroids compared to non-LAM fibroblasts. Treatment with sorafenib, a multikinase inhibitor, significantly reduced invasion, outperforming Rapamycin. We also confirmed TSC2-null AML cells as key VEGF-A secretors, which was suppressed by sorafenib in both AML cells and LAMFs. These findings highlight VEGF-A and bFGF as potential therapeutic targets and suggest multikinase inhibition as a promising strategy for LAM.

**One Sentence Summary:** Using 3D spheroids and spatial transcriptomics, we identified LAMFs and LECs as key contributors to LAM, with bFGF and VEGF-A as potential therapeutic targets

## INTRODUCTION

Lymphangioleiomyomatosis (LAM) is a progressive lung disease characterized by the formation of neoplastic lesions, consisting of tumorous smooth muscle-like cells and HMB-45 expressing epithelioid-like cells (LAM-cells). These lesions lead to cystic destruction of lung tissue and invasion of bronchioles, blood vessels, and lymphatic vessels, ultimately causing severe obstructive lung disease (*1, 2*). Despite advances in understanding the disease, therapeutic options for LAM remain limited, and it continues to be a debilitating condition primarily affecting women of childbearing age.

One of the major challenges in studying rare lung diseases like LAM is the lack of disease-relevant, patient-specific tissues and animal models that accurately reflect the human LAM phenotype (*3*). The cellular and molecular mechanisms driving the extensive cystic remodeling of pulmonary airspace and parenchyma in LAM are not well understood, resulting in a paucity of known molecular targets for therapeutic intervention. LAM can manifest sporadically or present as part of tuberous sclerosis, driven by somatic or germline mutations in the tuberous sclerosis complex (TSC1 or TSC2) genes (*4–10*). LAM lesions consist of aggregations of epithelioid cells expressing HMB-45 and smooth muscle α actin (SMΑA) expressing myofibroblast-like cells (*1, 3, 11–14*) and lymphatic endothelial cells (LECs) (*15–21*). Yet the impact of these cell types on disease pathogenesis remains poorly understood.

Lung resident fibroblasts, which have been implicated in other conditions such as interstitial lung disease, lung cancer, and kidney fibrosis (*22–25*), are believed to modulate the microenvironment in LAM, potentially activated in a manner similar to carcinoma-associated fibroblasts. Simultaneously, LECs are recognized as key contributors to the progression of neoplastic lesions, possibly facilitating the trafficking of LAM cells through the lymphatic system, thereby promoting metastasis (*26*). Several pathways have been suggested in the literature as contributors to LAM cell detachment and migration via the lymphatic system to other organs (*17, 27–30*).

Understanding the roles of LECs and LAMFs is critical for identifying new therapeutic targets and developing more effective treatments. However, the precise contributions of LECs to LAM pathogenesis, including their involvement in lymphangiogenesis and interaction with LAMFs, remain unclear. Most cellular models of LAM study cells in isolation within 2-dimensional (2D) culture environments, these are reductionist and often lose important properties like differentiation, polarization, cell-cell communication, and interactions with the extracellular matrix (ECM). At the same time wound healing, inflammatory processes, and hyperproliferation are artificially promoted (*31*). These limitations hinder our understanding of the intercellular interactions driving LAM pathogenesis, particularly between LAMFs and LECs.

Current therapeutic strategies predominantly target the core TSC2^-/-^ LAM cells, potentially overlooking the role of simultaneously activated cells, such as LAMFs and LECs, within LAM nodules. These overlooked components may play a crucial role in disease progression, underscoring the need for a more comprehensive approach that addresses the interactions between LAM-associated cells and their microenvironment. Rapamycin, an mTOR inhibitor, is the standard therapy for LAM patients with declining lung function (*32*). It inhibits mTORC1 activity, preventing the increased cell growth, proliferation, and survival associated with TSC2 mutations (*4, 30, 33–36*). Current guidelines recommend Rapamycin therapy for patients with an FEV1 <70% (*32*).

However, LAM shares mechanistic similarities with other neoplastic diseases, including cytokine secretion and the activation of pathways that promote cell growth and migration (*37–40*). Given this context, investigating multikinase inhibitors that target multiple cell types in the LAM microenvironment is crucial, especially in patients with advanced disease. sorafenib, an FDA-approved multikinase inhibitor, was chosen as a proof-of-concept therapy due to its known effects on tumor cell growth, angiogenesis, and metastasis (*41–43*). Sorafenib also inhibits eIF4E, a factor upregulated in LAM downstream of mTOR activation, and blocks VEGFR-2 and VEGFR-3, thereby potentially preventing LAM-associated lymphangiogenesis (*3, 43–45*). Additionally, sorafenib has shown promise in a mouse model of LAM, improving outcomes alongside Rapamycin (*46*). To better understand the interactions between LAMFs and LECs, we developed 3D spheroid models incorporating these cell types. Our study aimed to elucidate their collaborative roles in disease progression and evaluate the therapeutic potential of sorafenib in modulating these interactions and invasive behaviors.

## RESULTS

### In-situ LAM Nodule Formation confirms LEC Involvement

To validate LEC distribution and association with LAM-nodules we evaluated lung tissues from two end-stage LAM donor lungs (**Fig. 1 and table S1**). Hematoxylin and eosin (H&E) staining of these LAM lung tissues revealed a range of cyst sizes, indicative of alveolar simplification, a hallmark of LAM pathology (**Fig. 1A**, red arrows). Using Opal multiplexed staining, we assessed the co-localization of LAM-associated proteins within a single tissue, providing enhanced spatial insight into cell distribution within the LAM lung. LAM nodules (orange arrows), consisting of SMΑA expressing cells, were observed in close proximity to the cysts (white arrows, **Fig. 1B-F and fig. S1**). Higher magnification of these regions highlighted the presence of epithelioid-like, HMB-45 positive LAM cells within the SMΑA-expressing nodules (**Fig. 1C-F**, cyan). Additionally, VEGFR3 and PDPN-expressing LECs were clearly identified in and around these HMB-45/SMΑA-positive LAM nodules, particularly near cysts, as well as in spindle-shaped myofibroblast-like LAM cells (**Fig. 1B-F and fig. S1**). These observations align with the well-documented pathological features of LAM, as described in the literature (*2, 12, 20, 47, 48*). The presence of LECs invading and surrounding LAM nodules, especially as seen in Figure 1F, further supports their involvement in LAM pathogenesis, consistent with previous studies. The confirmation of LECs around LAM nodules reinforces their proposed role in the disease (*19–21*).

**Figure 1:**
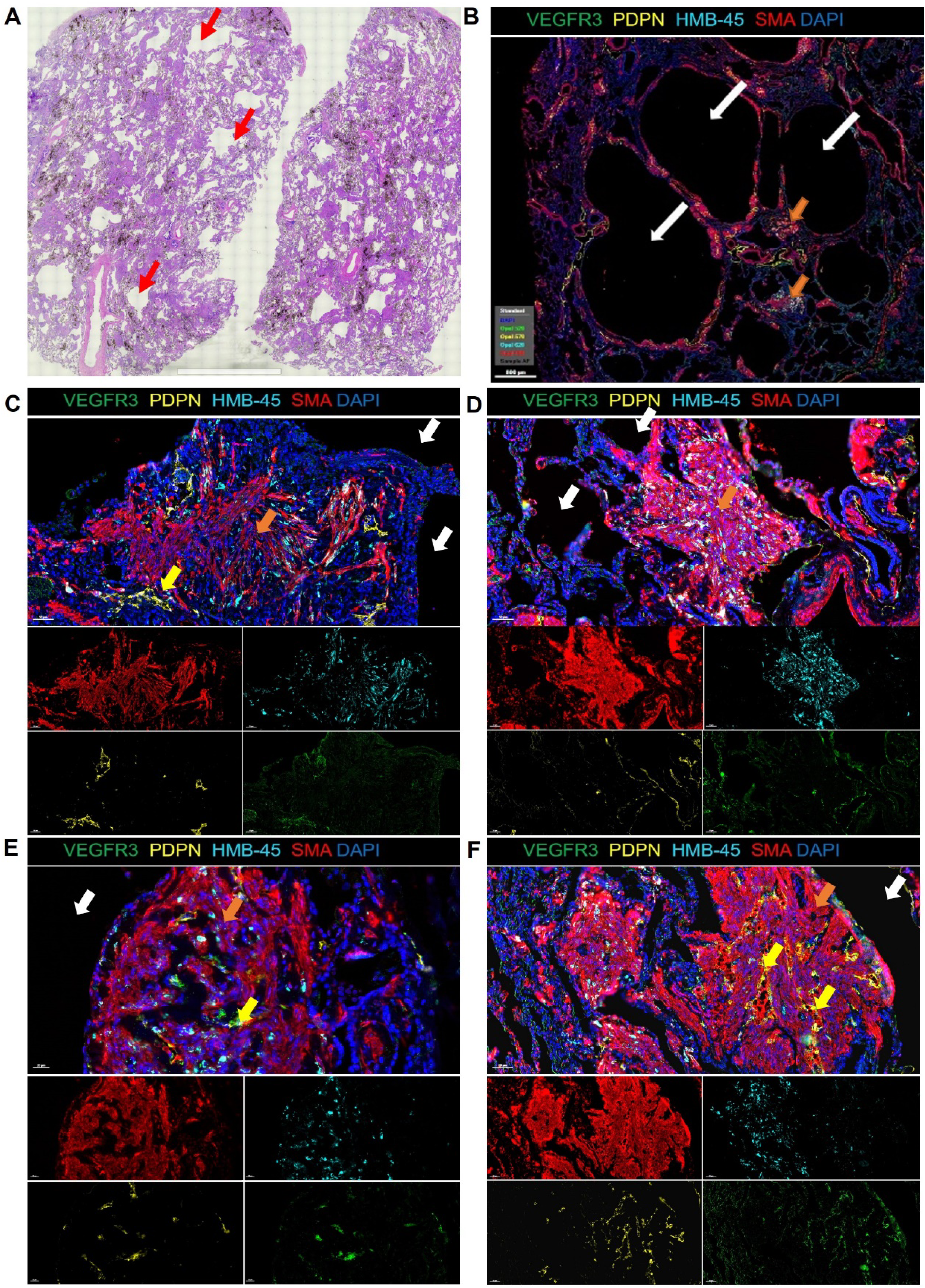
Identification of LAM nodules in patient LAM lung tissue. (A) Tiled 20x image of H&E staining of LAM lung tissue, red arrows indicate simplified alveoli. (B-F) Representative images of LAM lung tissues stained with VEGFR3 (green), PDPN (yellow), HMB-45 (cyan) and SMΑA (red). Nuclei are counterstained with DAPI (blue) in all images. White arrows indicate alveolar cysts, orange arrows highlight LAM nodules forming near cysts and yellow arrows in D indicate PDPN and VEGFR3 expressing LEC recruited to LAM nodules. Scale bars represent 2mm in A, 500 µm in B and 50 µm in C-F.

### Spatial Transcriptomic Analysis Identifies Key LAM Core Genes within the LAM Nodule Core

To further investigate the spatial organization of LAM cells, we conducted *in situ* spatial transcriptomics on two independent regions of formalin-fixed paraffin-embedded (FFPE)-prepared LAM donor lung tissue (**Fig. 2 and fig. S2-4**). Transcriptomics from 3343 and 2595 spots were obtained at a median depth of 8283 and 7536 UMIs/spot and 4317 and 4090 genes/spot for LAM_D1 and LAM_D2, respectively (**fig. S2**). Unbiased clustering and spot feature analysis classified the spots into ten and nine clusters for the integrated data of LAM_D1 (**Fig. 2A-C**) and LAM_D2 (**Fig. 2 E-F**), respectively. Based on both pathological identification of LAM nodules in the H&E-stained tissue sections (**Fig. 2A and E**) and expression of LAM-Core genes, such as ACTA2, ESR1, PMEL, RAMP1 and VEGFD (**databases S1-2**) (*49*), exhibited significantly higher expression levels in the LAM-Core cluster (**Fig. 2D and G**) were able to assign clusters 7 and 6 as LAM-Core clusters in LAM_D1 and LAM_D2, respectively.

**Figure 2:**
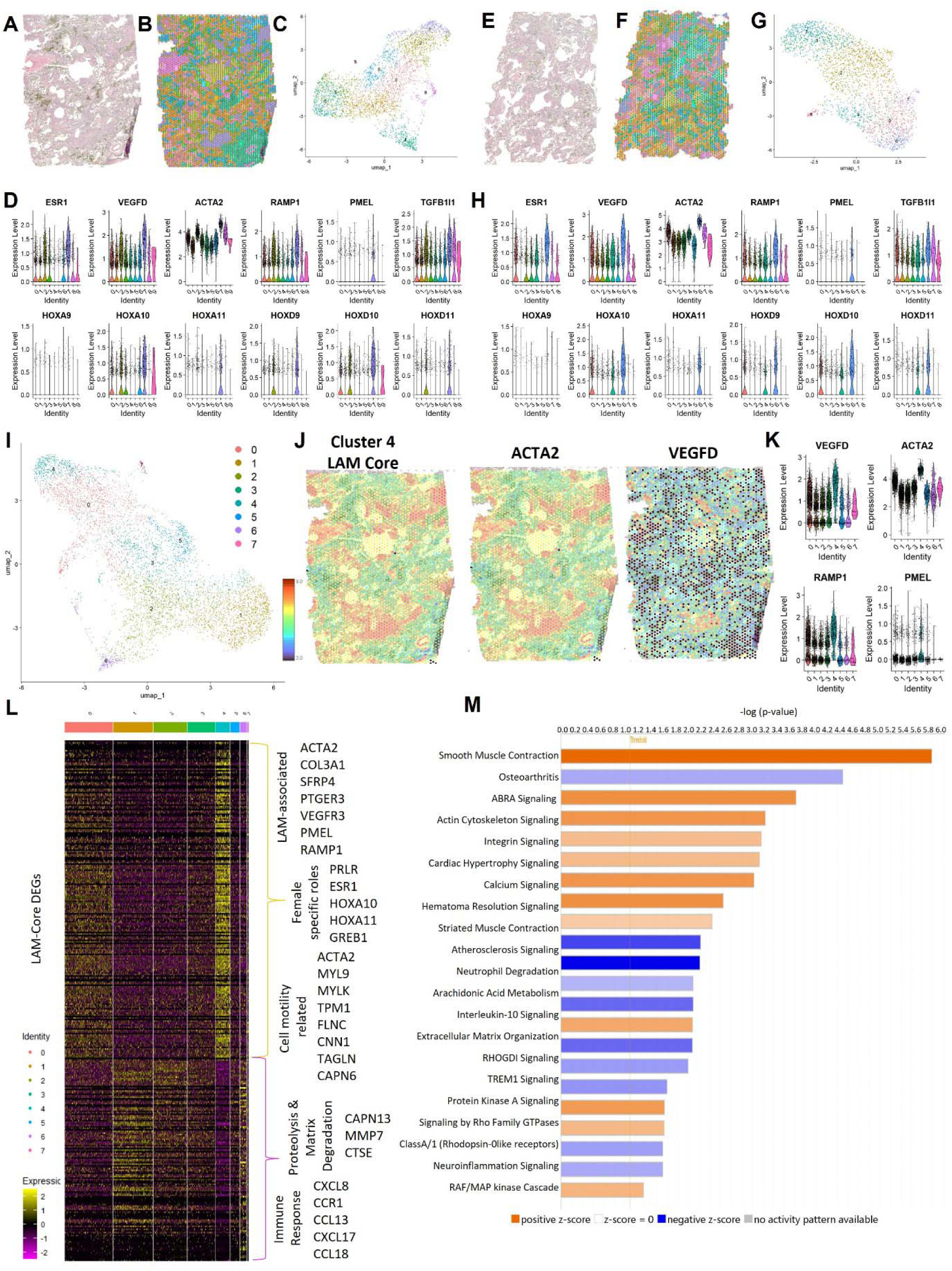
Spatial-omic analysis of LAM tissue highlights LAM-Core regions. (A & E) H&E staining of LAM lung tissues used for spatial transcriptomics. (B & F) Spatial mapping of the gene clusters for each of the lung tissues identified in the UMAPS provided in C & G, respectively, the colours represent the distinct cellular clusters categorized by gene expression on one of spot transcriptome. (D&H) Violin plots for established LAM-associated genes represented across each cell cluster for LAM_D1 and LAM_D2 lung tissues. (I) UMAP where colours represent the distinct cellular clusters categorized by gene expression on one of spot transcriptome for the combined dataset from both LAM tissues. (J) Spatial localization of well-established LAM genes ACTA2 and VEGFD from LAM_D1 lungs showing LAM nodule localization correlating to high gene expression. (K) Violin plots of LAM-Core genes in the combined dataset identifying cluster 4 as the LAM-Core. (L) Heat map for gene expression for the combined dataset where each column represents the colour coded cell cluster for all differentially expressed LAM-Core enriched genes with key up and down regulated genes highlighted. (M) Significant IPA canonical pathways enrichment in the LAM-Core. Orange represents positive z-scores, and blue represents negative z-scores.

For subsequent analysis of the transcriptomics data, we integrated both datasets together creating a new UMAP of cell clusters (**Fig. 2I**). Spatial mapping of the turquoise cluster (cluster 4) onto the H&E lung tissue section highlights its association with LAM nodules. We further demonstrate the localization of well-established LAM markers ACTA2 and VEGFD within this cluster (**Fig. 2J**), and the violin plots also include LAM core genes PMEL and RAMP1 (**Fig. 2K**). The heat map highlights all LAM-Core significant differentially expressed genes (DEGs) between the clusters (**Fig. 2L**). These genes include those well established in the pathogenesis of LAM in addition to TGFRB3, HOXA9-11, HOXD9-11, COL14A1, COL3A1, SFRP4, PTGER3 and VEGFR3 (**database S3**). Significantly upregulated genes also include genes that have been directly linked to female physiology and disease including prolactin receptor (PRLR), estrogen receptor 1 (ESR1), Hox family genes (HOXA10 and HOXA11) and growth regulation by estrogen in breast cancer 1 (GREB1) and many genes associated with processes related to muscle contraction, cytoskeletal organization and cellular motility including ACTA2, myosin light chain 9 (MYL9), myosin light chain kinase (MYLK), tropomyosin 1 (TPM1), filamin C (FLMC), transgelin (TAGLN) calponin 1 (CNN1) and calpain 6 (CAPN6). Significantly down regulated genes include genes related to proteolysis and matrix degradation including matrix metalloproteinase 7 (MMP7), cathepsin E (CTSE) and calpain 13 (CAPN13), likely to impact tissue remodeling and a number of genes related to immune response and inflammation including CXCL8 (IL-8), CCR1, CCL13, CXCL17 and CCL18 involved in chemotaxis and immune cell recruitment. To identify significantly enriched canonical pathways in the LAM-Core, we conducted Ingenuity Pathway Analysis (IPA) on the LAM-Core DEGs, applying a z-score threshold of 1 and p-value of 1.5 (**Fig. 2M**). The most significantly enriched pathways included those related to cell contraction and motility, calcium dynamics, extracellular matrix interactions and immune cell regulation. Additionally, several kinase signaling pathways were highly represented, including protein kinase A signaling, RAF/MAPK signaling, and Rho A kinase related signaling (**Fig. 2M, fig. S3C-D and database S4**). The LAM-Core significant DEG were also mapped for their distribution across the sorted cell type annotations based on human single cells dataset in the Lung Gene Expression Analysis (LEGA) and were found to be predominantly mesenchymal in origin (∼65%) with ∼15% endothelial, 17% epithelial and 3% immune cells **(fig. S3A)**. The proportion of endothelial related transcripts in the LAM-Core is supported by expression of LEC genes LYVE1, PDPN and SOX18 also in this same cell cluster (**fig. S3B**).

### Azimuth (HLCA) Analysis Reveals Myofibroblasts as Key Contributors to LAM Pathology

To assign lung cell types to the unique molecular identifiers (UMIs) of spatial-transcriptomics data we mapped our data to a comprehensive expression atlas of the human lung (50, 51). The cell clusters in the spatialomics data sets mapped to 10 different lung cell types in the Lung Map Atlas (**Fig. 3A-D and table S4**). LAM-Core genes in cluster 7 (purple, LAM_D1) and cluster 6 (blue, LAM_D2) (**Fig. 3A and C**, respectively), mapped to myofibroblasts, activated fibroblasts in the LAM-lung tissues, after robust cell-type decomposition (RCTD) using a single cell reference-based model (**Fig. 3B, D**). Mapping the spatial localization of the myofibroblast cluster we can see that it co-localizes with the pathologist-identified LAM-nodules, supporting the presence of LAM-Core cells in close association with a myofibroblast phenotype (**Fig. 3E-F**). Looking more closely at the relative expression of LAM-Core genes and LEC genes in violin plots we can see that these genes are most represented in cluster 7 for LAM-D1 (**Fig. 3G**) and in the myofibroblast-mapped population (**Fig. 3H**). To summarize, our spatial transcriptomics analysis identified LAM-Core genes within a cell cluster that spatially overlaps with pathological LAM nodules in lung tissue. The cells most closely associated with this region include myofibroblasts with endothelial and epithelial cells. The endothelial cells express lymphatic endothelial markers suggesting a close relationship with LECs in the LAM nodules. To further investigate the interactions between LAMFs and LECs we obtained primary LAM-patient derived fibroblasts and commercially available LECs to establish co-culture systems to evaluate cellular interactions.

**Figure 3:**
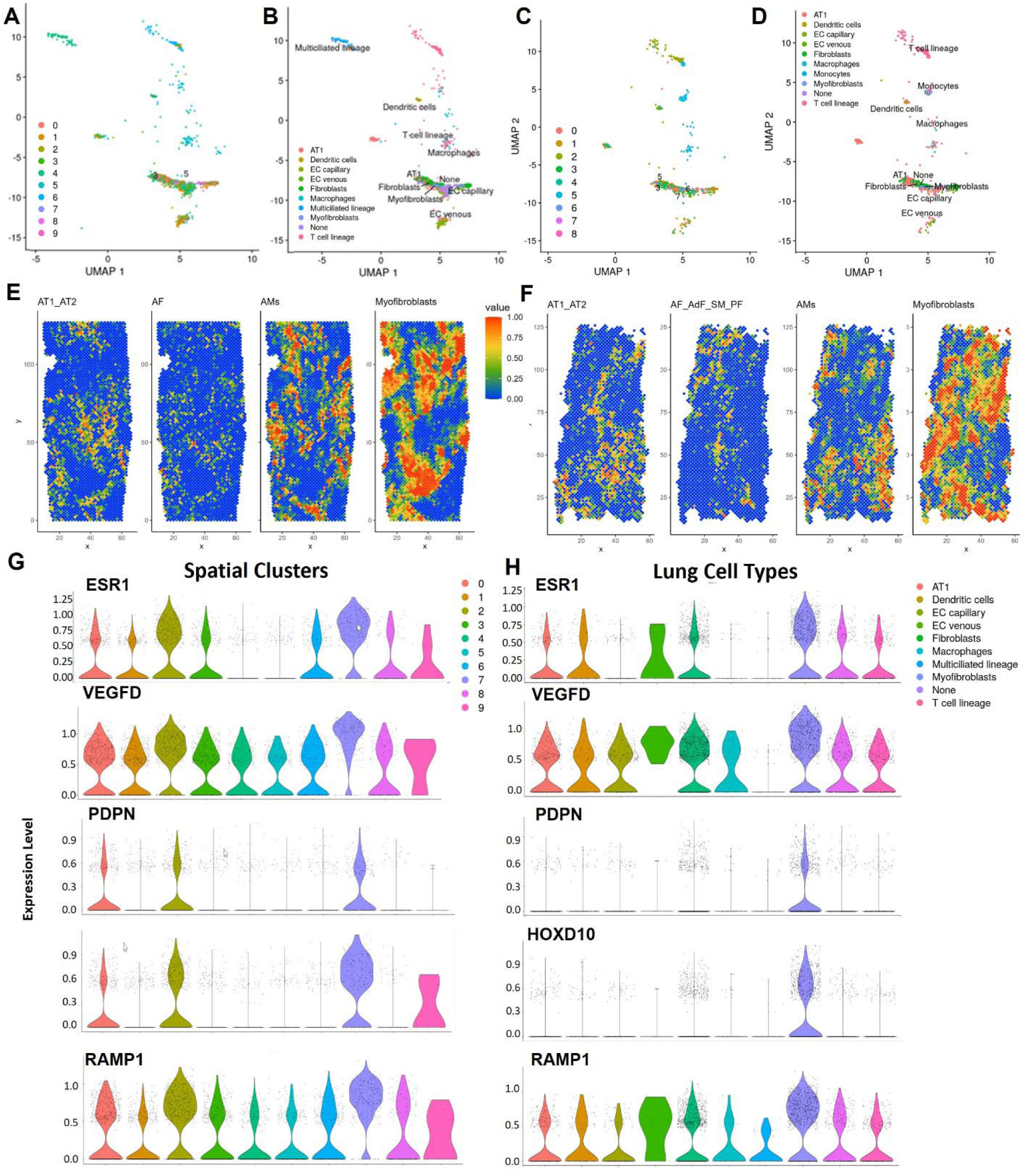
LAM-Core enriched gene signature maps to Myofibroblasts in Azimuth Lung v2 (HLCA) database. (A & C) Spatial transcriptomic gene clusters from LAM_D1 and LAM-D2 tissues, respectively mapped to Lung v2 dataset for level 3 annotation. (B & D) Spatial gene clusters represented by cell type signatures in human lung tissues for LAM_D1 and LAM_D2, respectively. (E & F) Robust cell-type decomposition (RCTD) images representing LAM_D1 and LAM_D2 cell-associated gene expression. (G & H) Violin Plots for LAM_D1 tissues showing relative gene expression of LAM-Core genes and LEC gene PDPN with highest expression in Cluster 7 (G) mapping to myofibroblasts (H).

### Activated LAMFs Exhibit Elevated PDGFRB, αSMAA, and mTOR and Increased Invasion

We obtained donor derived LAMF cells from Dr. Krymskaya (UPenn, **table S1**) and compared these to human lung fibroblasts (HLFs) from donors with no evidence of chronic lung disease (**table S1**). Despite some donor-to-donor variability, the LAMFs expressed significantly higher amounts of SMΑA protein (**Fig. 4A-B**). We also evaluated expression of mTOR and its downstream target P70S6K (**fig. S4A-B**). Although a significant increase in phosphorylated mTOR was noted in LAMF, the overall ratio of phosphorylated to unphosphorylated protein was not significantly changed (**fig. S4A**). Despite some donor variability, there was a significant increase in phosphorylated P70S6K and an increase in the ratio of phosphorylated to unphosphorylated protein in the LAMF, suggesting LAMF have changed mTOR activity (**fig. S4B**). To evaluate functional changes, focusing on matrix invasion, we generated matrix embedded spheroids. Cell number and media composition were optimized to reproducibly generate spheroids from both HLF and LAMF using aggrewell plates to generate spheroids consistent in size and cell number (**fig. S5A-B**). After 24h the spheroids were embedded in Matrigel and their growth and invasion into the surrounding matrix was quantified over a 7-day period (**Fig. 4B-E**). Physical properties of compactness, perimeter, and solidity were measured using an image analysis command pipeline generated in CellProfiler (Broad Institute, detailed in the **Online Data Supplement and fig. S5C-E**). Each mapped spheroid was checked manually, and manual spheroid edge identification was used where inaccuracies were observed due to low contrast differences between cells and background (**fig. S5**). LAMF spheroids had significantly greater compactness starting from Day 3 and sustained through Day 7 than HLF spheroids (**Fig. 4D**) indicating a highly irregular shape compared to HLF. A compact spheroid (circular) would have a value of 1 and irregular objects have a number greater than 1. LAMF also has significantly decreased solidity than HLF spheroids, again starting at Day 3 and sustained through Day 7 (**Fig. 4E**) indicating a higher convexity and irregular boundaries as well as being more porous. Interestingly, although trending, there was no significant difference in perimeter (data not shown). The phase contrast images provide representative examples of HLF spheroids and LAMF spheroids after 7 days of 3D culture, while LAMF are noted to be invading into the matrix, the HLF spheroids typically formed SMαAll nodules rather than projections into the ECM (**Fig. 4B-C**).

**Figure 4:**
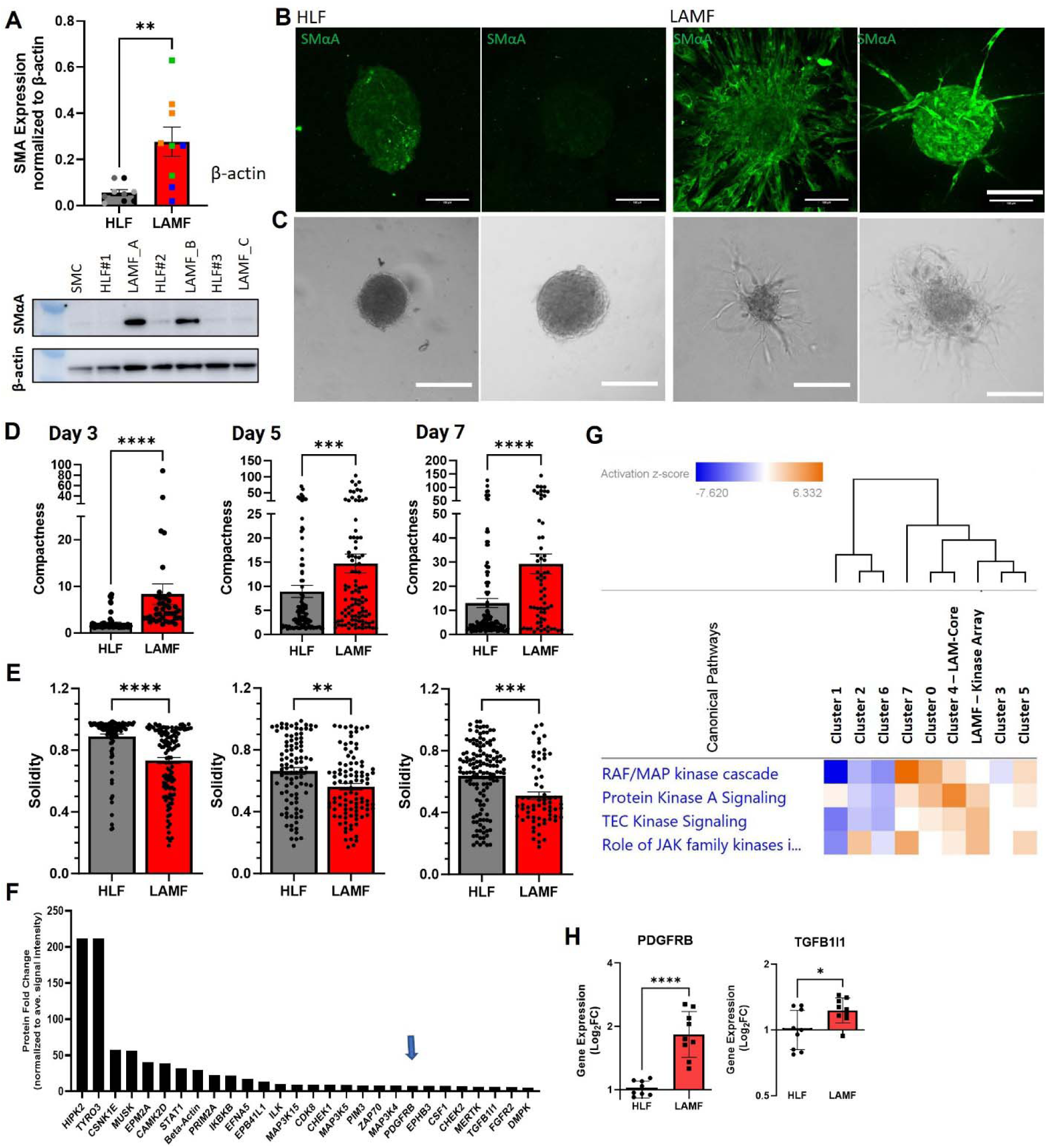
LAMFs represent an activated lung fibroblast phenotype compared to Human lung fibroblasts (HLFs). (A) Relative expression of SMαA comparing HLFs and LAMFs from 3 independent donors with a representative western blot, quantification normalized to β-Actin. (B – C) Representative SMαA (green, C) and phase contrast (D) images of spheroids generated from HLF and LAMF, scale bars = 100µm. (D - E) Quantification of changes in the compactness (D) and solidity (E) of spheroids over 7 days comparing LAMF to HLF. Each dot indicates a spheroid and a minimum of 10 (range 11-78) spheroids were evaluated. N=3 biological replicated per cell type. (F) Kinase array for LAMF representing expression in LAMF relative to the average signal intensity across all proteins evaluated. (G) Heat map of canonical pathways comparing the integrated spatial transcriptomics data with the kinase array data showing kinase related pathways data shown has a cut off z-score >1 and Log_10_P value >1.5, orange is higher pathway activation and blue is pathway inhibition. (H) qRT-PCR comparing gene expression of PDGFRB and TGFB1I1 in HLF and LAMF. Data represents mean ± S.E.M. with significance represented by * p<0.05, ** p<0.01, **** p<0.0001.

As multiple pathways regulating kinases, cell adhesion and ECM-interactions were confirmed to be transcriptionally upregulated in the LAM-Core, we evaluated the expression of a panel of kinases in isolated LAMF through a kinase array (**Fig. 4F and database S5**). The data shown in **Figure 4F** highlights the kinases with the highest expression levels in LAMF; the proteins that had a greater than a 1.25-fold change increase compared to the median protein level of all kinases are featured. The highest expressed protein kinases were HIPK2 and TYRO3 (**Fig. 4F**). However, other notable highly expressed proteins included PDGFRB, MAPK10, MAK4K5, FGFR1, CAMKIID and TGFB1I1 (TGF beta 1 interacting protein 1). The heatmap in **Figure 4G** compares the changes in canonical pathways focused on kinase-related signaling pathways including the integrated transcriptomic data and kinase array data. Hierarchal clustering shows close association between the kinase expression in the transcriptomics data for cluster 4 (LAM-Core) and the protein kinase array. We confirmed the gene expression levels of PDGFRB and TGFB1I1 in three independent donor-derived lines of LAMFs to HLFs and confirmed significantly elevated levels of these genes in LAMFs (**Fig. 4H**). This data supports a significant change in fibroblast phenotype in LAM lung tissues.

### Multicellular Spheroid Models Reveal Altered Fibroblast-Endothelial Cell Interactions in LAM

The multiplexed imaging and spatial transcriptomics data highlights close proximity of LEC and activated fibroblasts in the LAM nodules. To begin to understand the complexity of cellular interactions between LAM-cells, LAMFs and LECs in LAM pathogenesis we next established methodology to reproducibly generate multicellular spheroids comprising of LAMF and LEC (**Fig. 5**). Using PROX1, SOX18, LYVE1 and PDPN as a panel of core LEC genes we spatially mapped LEC into the lung tissue (**Fig. 5A**). Overlay of these genes with a signature of 5 LAM-Core genes (RAMP1, ACTA2, PMEL, VEGFD and HOXD10) indicates a close association of both cell types spatially in the LAM tissues (**Fig. 5B**) and transcriptionally these gene signatures are enriched in the LAM-Core cluster (**Fig. 5C**, blue). A 3D multicellular model containing both LAMF/HLF and LEC was generated to evaluate inter-cellular interactions in LAM pathogenesis (described in the **Online Data Supplement** and **fig. S6A-C**). Commercially available lung microvascular lymphatic endothelial cells (HpMVLEC, referred herein as LECs) were evaluated for the expression of LEC markers including expression of PDPN, PROX1 PECAM and VE-Cadherin (CD144). PDPN expressing lymphatic endothelial cells comprised of 96.5% of the endothelial cells by FACS and IF staining, validating the manufacturer’s description (data not shown). After 24 hours of co-culture, the spheroids comprised of a green-labelled LAMF core surrounded by red-labelled LEC (**Fig. 5D and fig. S6D**). The cellular distribution was confirmed through staining for SMαA for the LAM-cells and PDPN for the LECs (**fig. S6B**). After 3 days of spheroid co-culture there were minimal differences between the HLF and LAMF co-cultures spheroids (**Fig. 5E**). However, over a 7-day evaluation period the LAMF-LEC spheroids had significantly higher matrix invasion (**Fig. 5F-H**) and significantly altered spheroid physical parameters (**Fig. 5G**). The LAMF-LEC spheroids had a significant and substantial increase in spheroid compactness and perimeter, and decrease in solidity, reflecting increased invasion compared to HLF-LEC controls (**Fig. 5G-H**). Evaluation of live cell images indicate that most of the invading edges of the LAMF-LEC spheroids comprise of both cell types (**Fig. 5F and fig. S6E**).

**Figure 5:**
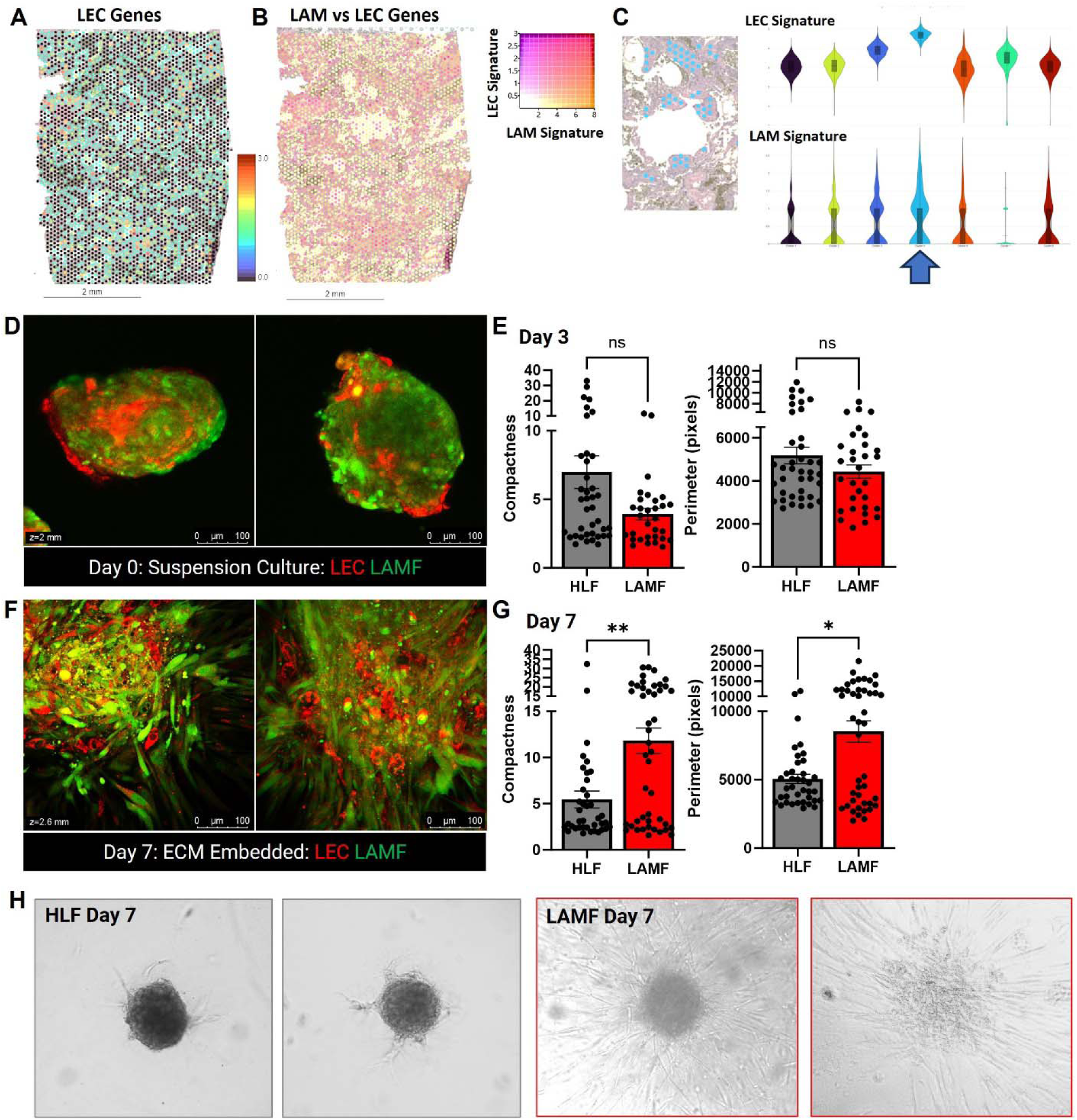
LAMF-LEC organoids have increased invasion into the ECM. (A) Spatial heatmap of localization of LEC genes in LAM_D1 tissue (SOX18, PDPN, LYVE1 and VEGFR3). (B) Co-localization of core LEC gene signature and LAM-Core signature genes in LAM lung tissue LAM_D1. (C) Violin plots showing highest expression of both LEC signature genes and LAM-Core signature genes in blue cluster 4 which spatially maps to histological regions of the lung tissue representing LAM nodules. (D) Representative IF images of LAMF-LEC spheroids with CellTracker-Red labelled LEC and CellTracker-Green labelled LAMF 24 hours after seeding in 3-D culture conditions (E) Quantification of changes in the compactness perimeter and solidity of the co-cultured spheroids over 3days comparing LAMF to HLF. Each dot indicates a spheroid and a minimum of 10 (range 11-78) spheroids were evaluated. (F) Representative images of LAMF-LEC spheroids embedded in ECM after 7 days. (G) Quantification of changes in the compactness perimeter and solidity of the co-cultured spheroids over 7 days comparing LAMF to HLF. Each dot indicates a spheroid and a minimum of 10 (range 11-78) spheroids were evaluated. (H) Representative phase contrast images of HLF and LAMF after 7 days of 3-D culture. In all experiments N=9 experimental repeats. Scale bars represent 100 µm. Data shown represents mean±SEM., significance represented by *p<0.05, **p<0.01 and ***p<0.001.

### sorafenib Inhibits Matrix Invasion of LAMF Spheroids and LAMF-LEC Multicellular Spheroids More Efficiently than Rapamycin

LAM cells are known to activate PI3K/AKT and ERK/MAP kinase signalling (*34*), in addition to possible induction of angiogenesis through soluble factors such as VEGF-A, PDGF and VEGF-D targeting VEGFR2, VEGFR3 and PDGFR-mediated signalling in the LAM microenvironment (*48, 52–54*). The data presented in this manuscript highlights the multicellular involvement in LAM pathogenesis and the augmentation of multiple kinase pathways in the LAM-Core. We therefore selected an FDA-approved multikinase-inhibitor, sorafenib, to target multiple cell types and signalling pathways to prove the importance of multicellular models and the need for combined signalling pathway targeting in the development of therapeutics for LAM. We compared sorafenib to rapamycin, an mTOR inhibitor that is the current and only approved therapeutic to slow the progression of LAM (33) on LAMF spheroids and LAMF-LEC multicellular spheroids (**Figs. 6-7**). sorafenib inhibits VEGFR2-, VEGFR3-, and PDGFRβ-mediated signaling targeting the MAPK/ERK pathway to prevent phosphorylation of downstream targets eIF4E and MEK1/2 and ERK1/2 (*42–44*). To evaluate the effect of sorafenib on the physical properties of the LAMF and HLF spheroids we imaged vehicle-treated (Veh), rapamycin-treated (Rapa) and sorafenib-treated (Sora) daily from day 3 to day 7 of 3D culture. Image analysis revealed an almost complete cessation of invasion in LAMF spheroids at 7µM sorafenib, as shown in **Figure 6**. While a small, but significant decrease in perimeter (**Fig. 6A**) was noted for Sora-treated HLF, there were no significant changes in spheroid properties in response to treatment with either Rapa or Sora after 3 days of culture (**Fig. 6A-C**). In LAMF spheroids, Sora significantly reduced both perimeter (**Fig. 6D**) and compactness (**Fig. 6E**) and increased solidity (**Fig. 6F**), while Rapa had no significant impact on LAMF at day 3 (**Fig 6D-F**). By day 7 Rapa and Sora both significantly and comparably reduced perimeter and compactness and increased solidity in the control HLF spheroids. Interestingly, Rapa had no effect on LAMF perimeter or compactness, while Sora significantly and substantially reduced both the perimeter and compactness to levels closer to that of vehicle treated HLF (**Fig. 6A-F**). The phase contrast images in **Figure 6G** are representative of the changes described above and clearly show the reduction in invasion of LAMF. Interestingly, a dose dependent decrease in cell metabolism was noted in sorafenib-treated LAMF at doses above 10 nM compared to HLF which only showed changes at 10 µM, indicating decreased cell viability and increased cytotoxicity specific to LAMFs (**Fig. 6G**).

**Figure 6:**
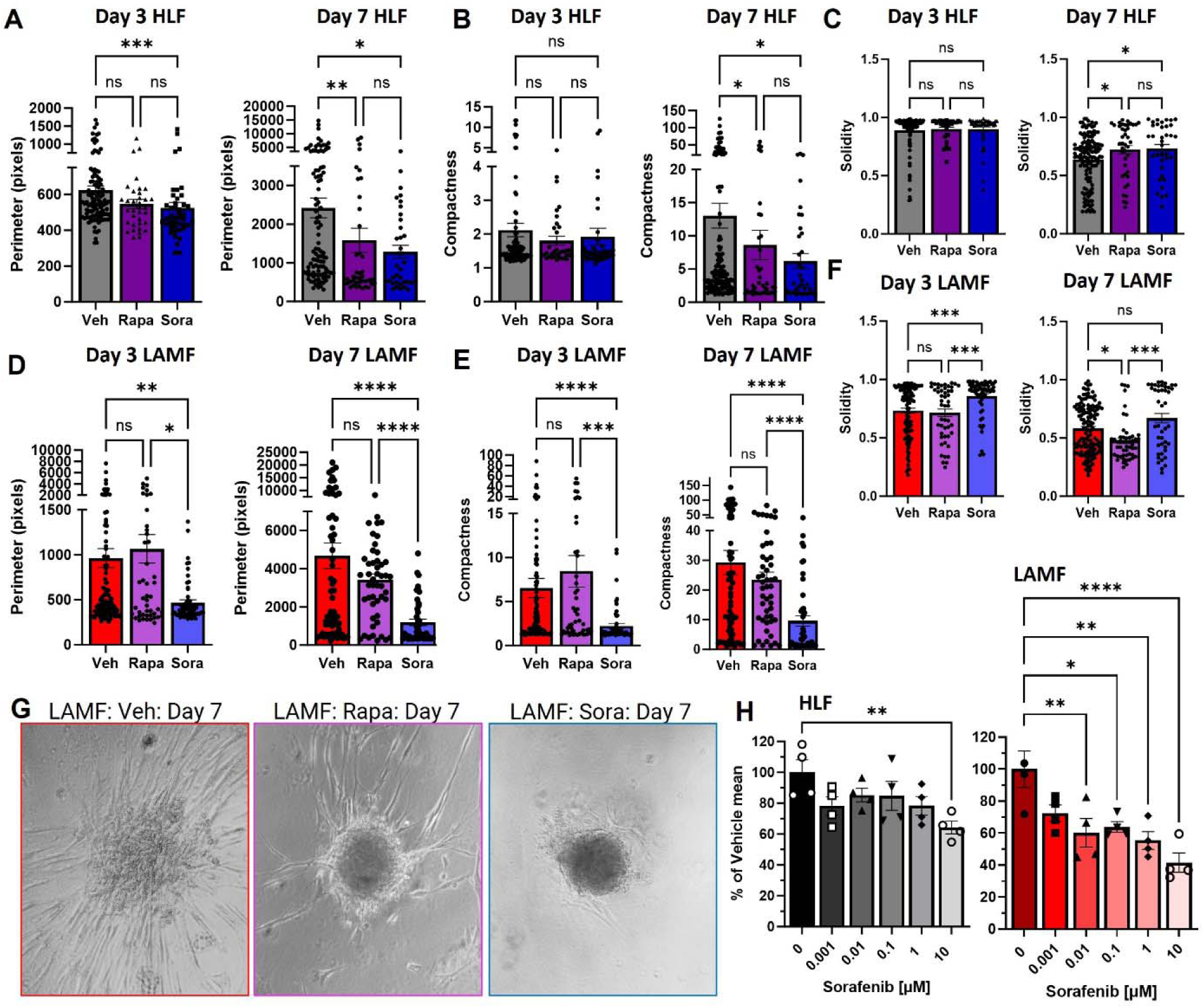
sorafenib treatment inhibits invasion of LAMF spheroids. (A-C) Changes in perimeter (A & D), Compactness (B & E) and Solidity (C & F) comparing day 3 and day 7 for HLF and LAMF spheroids treated with either 20 nM rapamycin (Rapa) or 7 µM sorafenib (Sora), compared to vehicle (Veh). Each dot indicates a spheroid and a minimum of 10 (range 11-78) spheroids were evaluated for each of 3 independent donors (N=3). (G) Representative phase contrast images of spheroids at Day 7 of treatment for LAMF comparing vehicle to Rapa and Sora treatments. (H) Presto Blue viability assays for HLF and LAMF in response to increasing doses of sorafenib, data is expressed as a percentage of the vehicle mean from three experimental repeats. Data shown represents mean±SEM, significance is represented by *p<0.05, **p<0.01, ***p<0.001 and ****p<0.0001, for N=3 independent donor cells.

**Figure 7:**
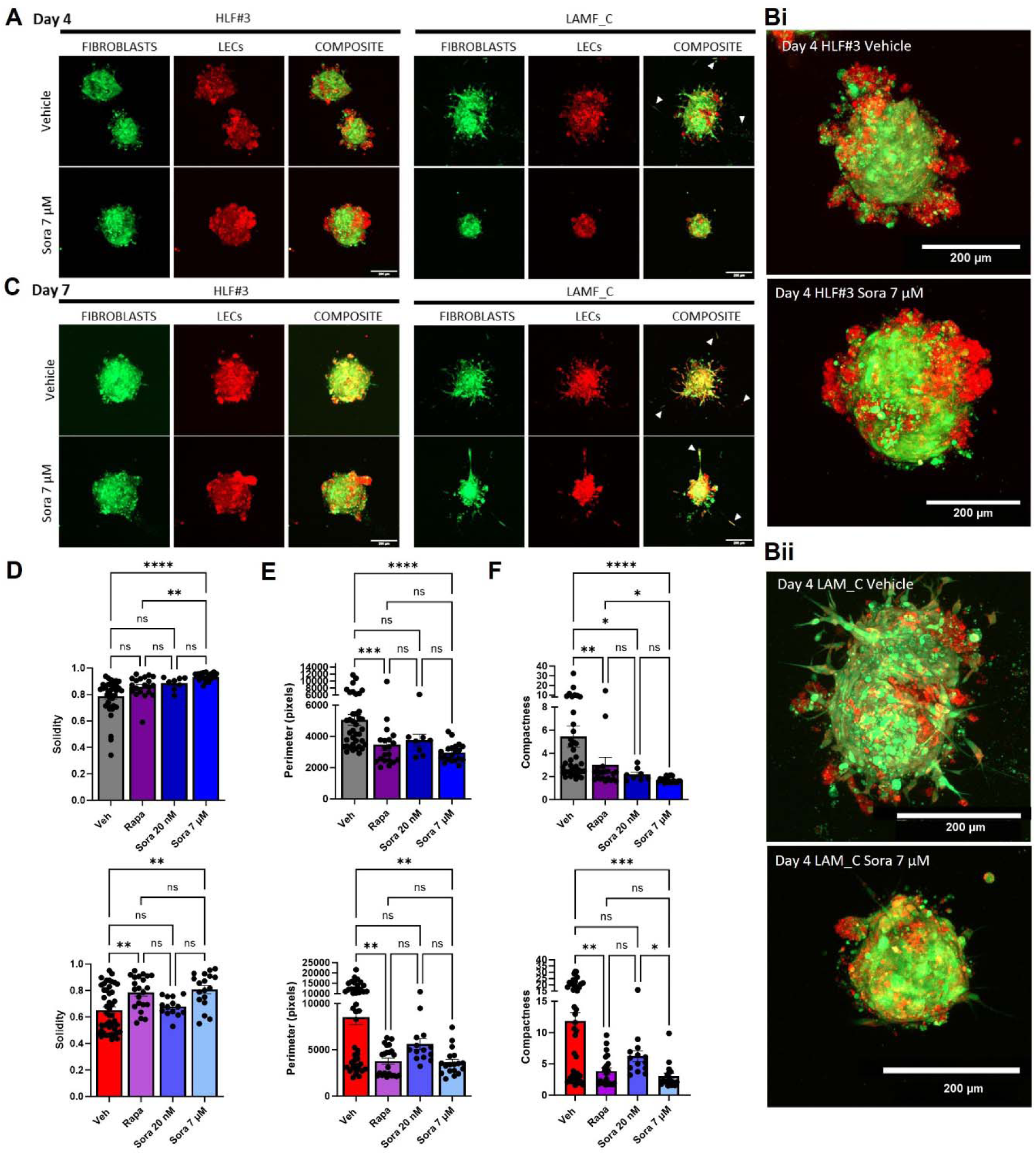
sorafenib treatment inhibits migration and invasion of LAMF-LEC spheroids. (A-C) Representative confocal images of spheroids at day 4 (A-B) and day 7 (C) of treatment with 7 µM sorafenib (Sora) or vehicle comparing HLFs and LAMFs co-cultured with LECs. Fibroblasts are stained with CellTracker Green and LECs with Cell Tracker Red. Scale bars in all images are 200 µM. D-F) Changes in Solidity (D), perimeter (E) and Compactness (F) at day 7 for HLF-LEC and LAMF-LEC spheroids treated with either 20 nM rapamycin (Rapa) or 20 nM or 7 µM sorafenib (Sora), compared to vehicle (Veh). Each dot indicates a spheroid and a minimum of 10 (range 11-78) spheroids were evaluated per donor. Data shown represents mean±SEM, significance is represented by *p<0.05, **p<0.01, ***p<0.001 and ****p<0.0001 for each of N=3 donors.

To determine whether the presence of LEC impacts the activity of sorafenib on the physical properties of the co-cultured LAMF and HLF spheroids we monitored Veh, Rapa and Sora treated spheroids daily from day 3 to day 7 of 3D culture (**Fig. 7A-C**). As shown in the representative images the HLF co-culture spheroid morphology does not change considerably in the presence of Sora, with the LEC (red) clustering on the outer edges of the HLF spheroids at day 4 and similar spheroid organization and size is observed at day 7 (**Fig. 7A-C**). On the other hand, LEC co-cultured with LAMF spheroids have notable matrix invasion at day 4 (**Fig. 7A-B**) which is almost completely inhibited in the presence of Sora at both days 4 and 7 of differentiation (**Fig. 7A-C**). Image analysis revealed an almost complete cessation of invasion in the LEC-LAMF spheroids at 7µM Sora. Unlike the monocultures, Sora had no significant impact on the co-cultured spheroids after 3 days for either the HLF or LAMF, suggesting some crosstalk between LECs and lung fibroblasts (**Fig. S6F-G**). After 7 days Rapa and Sora treatment significantly reduced perimeter and compactness with Sora increasing solidity in both HLF and LAMF co-cultured spheroids (**Fig. 7D-F**). It is worth noting that the Sora-treated LAMF have visual and physical properties after Sora treatment that are akin to LEC-HLF spheroids in vehicle (**Fig. 7B**).

### Intercellular Signaling Drives Changes in Cellular Phenotype Associated with Increase of Secreted VEGF-A and bFGF from TSC2^-/-^ LAM-Cells

It is not currently known how LECs are recruited to LAM nodules and whether their tissue invasion is sensitive to changes in LAMF. Considering that the main alteration and potential priming of both LAMF and LECs are driven by TSC2^-/-^ LAM cells, we evaluated changes in secreted growth factors and cytokines from both LAMFs and TSC-2^-/-^ cells. TSC-2^-/-^ (S102), and their gene corrected controls (S103) were derived from TSC2-deficient renal angiomyolipoma cells (AML) and kindly shared by Dr. Henske (Harvard, Boston). TSC-2^-/-^ cells have been shown to activate LAMF, upregulating growth factors, such as FGF7, and subsequently LAMF can generate soluble factors potentially influencing alveolar cell phenotype (*55*). To propose a mechanistic association with the recruitment of LEC to the LAM nodules we used an angiogenesis multiplexed ELISA panel to determine changes in secreted growth factors and cytokines from both HLFs, LAMFs, S102 and S103 cells in the presence of vehicle, rapamycin or sorafenib (**Fig. 8**). Although trending to increased levels for both VEGF-A and VEGF-C, there were no significant differences in the secreted factors from HLFs and LAMFs in the presence or absence of the inhibitors (**Fig. 8A**). This may be a result of a loss of phenotype with cell passaging *in vitro*. However, the secretions from the fibroblasts were 30-60-fold lower than those observed from the AML cells indicating that the major changes in secreted factors in the LAM-Core are more likely to come from the LAM-cells and not the LAMFs. Secretion of VEGF-A, VEGF-C and bFGF were significantly different between S102 and S103 and in response to both rapamycin and sorafenib treatment (**Fig. 8B**). Our results show that secretion of VEGF-A, a highly angiogenic growth factor, was significantly augmented in TSC2^-/-^ cells compared to TSC2 -expressing cells and both rapamycin and sorafenib significantly reduced its secretion in both cell types (**Fig. 8B**). Interestingly, both VEGF-C and FGF were secreted at significantly higher levels in the TSC-2 expressing cells, with rapamycin and sorafenib completely blocking secretion of VEGF-C in both cell types. A differential effect of rapamycin and sorafenib was observed on the TSC-2 expressing and deficient cells with rapamycin inhibiting or not changing secretion of bFGF and sorafenib significantly increasing secretion in TSC-2 expressing cells but significantly inhibiting secretion in TSC-2^-/-^ cells. Conditioned media from TSC-2^-/-^ cells significantly upregulated gene expression in normal human lung fibroblasts associated with activation and phenotypic switching to a myofibroblast (**Fig. 8C**). This data highlights a complex multicellular interaction, responsive to rapamycin and multikinase inhibition and highlights both 1) the necessity of multicellular systems to study LAM pathogenesis and 2) significant similarity of LAM-nodules to the interactions of activated fibroblasts and angiogenesis mechanism in cancer progression (*56*). **Figure 8D** highlights the significance of the model and mechanism presented in this manuscript and their potential to driving therapeutic innovation in LAM treatment.

**Figure 8:**
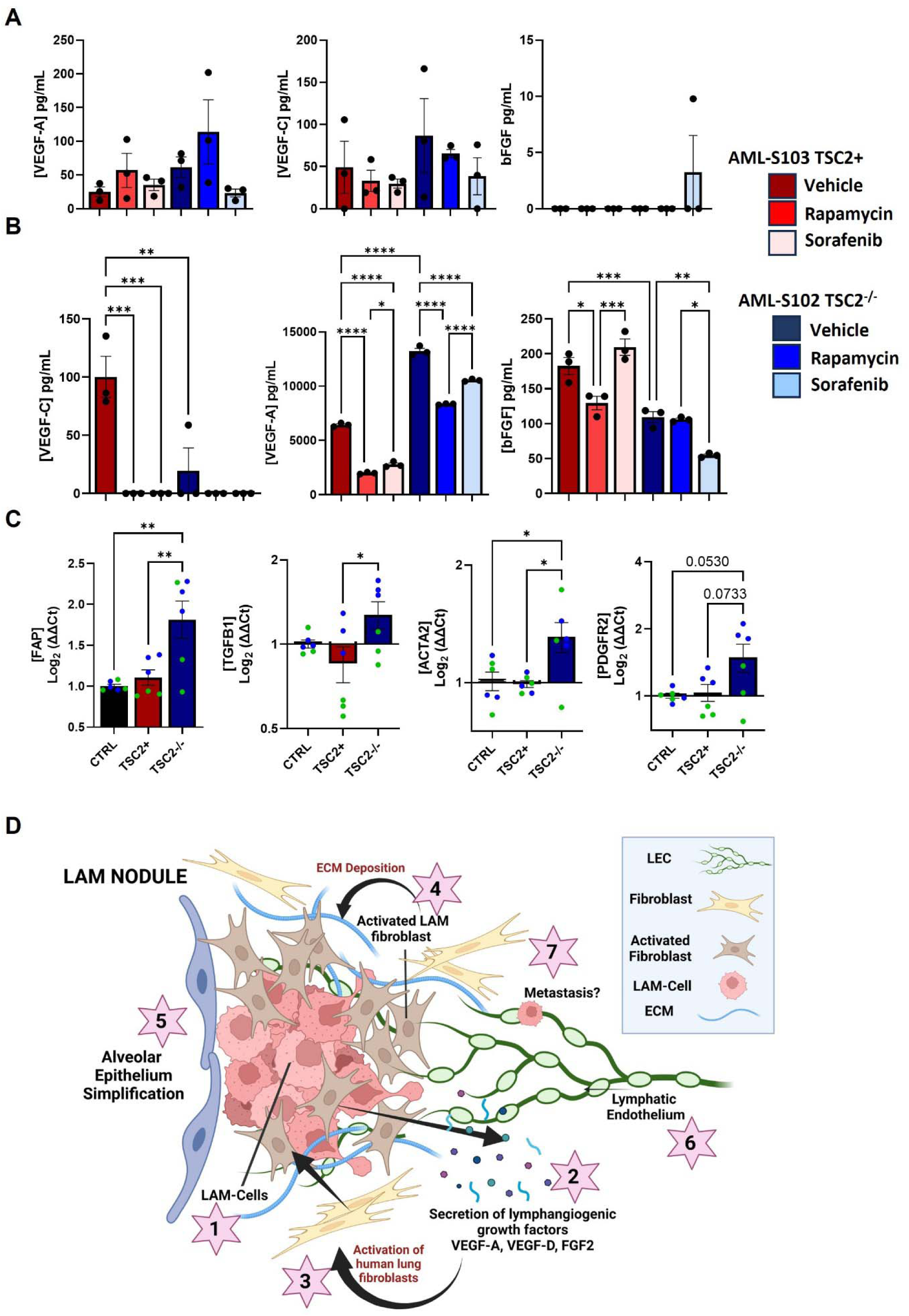
sorafenib inhibits secretion of pro-angiogenic cytokines from LAMF and AML-TSC2 cells. (A-B) secreted VEGF-A, VEGF-C and bFGF from HLF (red) or LAMF (Blue) (A) and AML-S102 (red) and S103 (blue) cells (B). (C) Gene expression of activated fibroblast markers, FAP, TGFB1, ACTA2 and PDGFRA, in HLF induced by supernatants from either HLF (black, control) or from AML-S103 (red, TSC2+) and S102 (blue, TSC2-/-) cells. Data shown represents mean±SEM, significance is represented by *p<0.05, **p<0.01, ***p<0.001 and ****p<0.0001 for each of N=3 donors. (D) Schematic depicting the proposed cellular interactions leading to LAM pathogenesis. LAM-cells (1, TSC2^-/-^) secrete high levels of VEGF-A, VEGF-D and FGF2 (2) which contribute to the activation of resident lung fibroblasts, generating activated myofibroblasts (3). Activated fibroblasts can lead to dynamic changes in cellular motility and influence the composition of the extracellular matrix (4), creating a unique LAM nodule niche. Activated fibroblasts can also influence alveolar stem cell behaviour (5) and these same secreted factors can recruit lymphatic endothelial cells (6) to LAM nodules, which may also provide a pathway for LAM metastasis to other organs (7).

## DISCUSSION

Our study highlights that LAM is a disease driven by complex interactions among multiple cell types, necessitating advanced multicellular models to fully elucidate its pathogenesis and to identify new therapeutic targets. The involvement of multiple signaling pathways further complicates LAM, suggesting that a multifaceted therapeutic approach will be essential to halt disease progression. LAM, as a slowly progressive and metastasizing neoplasm, falls within the family of perivascular epithelioid cell tumors (PEcoma). Although the exact origin of the cells that give rise to LAM nodules remains unidentified, these nodules are known to comprise various cell types, including TSC2-null smooth muscle-like cells, HMB-45-positive epithelioid-like cells (LAM cells), LECs, and activated fibroblasts (LAMFs), which closely resemble carcinoma-associated fibroblasts (CAFs).

Our spatial transcriptomics analysis confirms the presence of a set of LAM-Core genes, such as ACTA2, TGFBR3, HOXA11, HOXD11, COL14A1, and COL3A1, previously identified through single-cell RNA sequencing as being spatially localized within LAM nodules in patient-derived lung tissues (*49*). By mapping these LAM-Core cells to previously published lung datasets, we found that these cells are most closely associated with myofibroblasts. Myofibroblasts, as activated fibroblasts, play a crucial role in altering the cellular microenvironment by secreting extracellular matrix components, growth factors, and cytokines, which often facilitate tumor growth and progression (*57–60*). We show that normal HLFs respond to factors secreted by TSC2^-/-^ cells (analogous to LAM cells) by upregulating markers characteristic of activated fibroblasts. This finding supports the notion that while bona fide LAM cells may initiate the disease, their interactions with fibroblasts likely drive core pathological changes within the cellular niche, thereby perpetuating disease progression. Indeed, a population of alveolar fibroblasts (AF2s), which also resemble activated fibroblasts, is known to modulate alveolar cell phenotype, leading to alveolar simplification (*61*).

In addition, our study highlights the significant interaction between LAM cells and LECs, which migrate into LAM nodules. The changes in the cellular microenvironment, induced by both LAM cells and LAMFs, promote lymphangiogenesis by altering LEC interactions. Notably, we observed significant increase in VEGF-A secretion, a well-known driver of angiogenesis. Augmented VEGF-A expression is strongly linked to interstitial lung diseases (ILDs) and fibroblast activation, but its role in LAM’s tissue homeostasis and microenvironment regulation is not well understood. Nonetheless, the loss of TSC2 and subsequent mTOR upregulation, along with increased VEGF-A secretion, has been described previously, supporting our observations in LAM (*62*). Further investigation into the LAM cell secretome and its impact on human lung fibroblasts, LECs, and alveolar type 2 (AT2) cells will likely enhance our mechanistic understanding of the pathological signaling driving LAM, beyond the well-studied mTOR hyperactivation.

Our study underscores that LAM nodules resemble tumor nodules, driven by complex multicellular interactions and multiple signaling pathways. Notably, we demonstrate the utility of mono- and co-cultured spheroids to investigate the interaction of LAMFs and LECs, revealing a higher invasive capacity in these cells compared to normal fibroblasts and their co-cultures. This increased invasiveness is associated with significant upregulation of key signaling molecules, including VEGF-A, VEGFR2 and bFGF, suggesting a pro-tumorigenic mechanism within the LAM nodule microenvironment. Tumors with significant heterogeneity, including LAM nodules, often express PDGFRB and have been shown to attract LECs through the PDGF/PDGFRB axis (*63*) reinforcing the pathway for LAM cell-induced lymphangiogenesis and potential metastasis through lymphatic circulation (*16, 17, 52*). Additionally, the pro-neoplastic role of PDGF is further supported by its contribution to collagen formation, which may increase remodeling of the LAM microenvironment (*64*).

Furthermore, our study highlights the potential of multikinase inhibitors, such as sorafenib, in targeting these signaling pathways. Rapamycin, the primary drug of choice according to treatment guidelines, is currently only available for patients with reduced lung function in progressed LAM and while most of these patients do respond well, resulting in stabilization of lung function, it is not effective for all (*35, 65–68*). Rapamycin is also known to be primarily cytostatic and not cytotoxic, limiting its capacity for inhibition of tumorous growth and highlighting the need to identify new therapeutic targets for the treatment of LAM (*66–69*). sorafenib is an FDA-approved multikinase inhibitor (*41–43*) that is known to inhibit RAF/MEK/ERK signalling, leading to increased tumor cell apoptosis, decreased micro-vessel density, and reduced metastatic shedding of tumor cells (*41–44*) and inhibiting the phosphorylation of eukaryotic translation initiation factor (eIF4E), which is typically upregulated in LAM, downstream of the upregulated mTOR pathway (*3, 43–45*). In our model, sorafenib demonstrated superior efficacy compared to Rapamycin in suppressing LAMF invasion into the surrounding extracellular matrix (ECM), indicating its potential as a therapeutic option in advanced LAM by simultaneously targeting LAMFs and LECs. Previous studies have shown that sorafenib and other multikinase inhibitors can effectively impede tumor progression and reduce the likelihood of recurrence and metastasis (*41–43*). Our findings suggest that a combination of low-dose sorafenib with mTOR inhibitors, or even low-dose sorafenib monotherapy, may represent a promising therapeutic strategy for managing end-stage LAM, particularly in patients with high VEGF-D levels who are unresponsive to rapamycin. VEGF-D, currently utilized as a diagnostic biomarker for LAM with a serum cutoff value of approximately 800–1000 pg/mL, has shown a significant correlation with disease severity (*40, 69–72*). Additionally, lower diffusion capacity of the lung for carbon monoxide (DLCO) scores, which serve as functional markers of interstitial involvement and reflect tissue-level changes such as fibroblast activation and lymphatic disruption, have been associated with higher mortality in interstitial lung disease (ILD) (*73–75*). To enhance LAM management, refining VEGF-D cutoff levels through clinical studies or developing a comprehensive clinical scoring system that incorporates DLCO, FEV1, VEGF-D levels, and the assessment of lymphatic abnormalities could improve patient stratification and therapy effectiveness. Such an approach could be instrumental in assessing the efficacy of therapies like multikinase inhibitors. Ongoing clinical studies on Nintedanib, a multikinase inhibitor targeting FGFR, PDGFR, VEGFR, and TGF pathways, have shown potential in stabilizing fibrotic lung diseases like idiopathic pulmonary fibrosis, but its effectiveness in LAM remains under investigation. This highlights the importance of exploring a broader spectrum of kinase inhibitors for effectively halting LAM progress rapamycin, suggesting a more potent action beyond mere cytostatic activity. Notably, phase II clinical trials of nintedanib in idiopathic pulmonary fibrosis have shown promise in stabilizing disease progression, further supporting the potential of targeting a wider range of kinases in treating LAM (*76, 77*).

Looking ahead, focused research on the LAM cell secretome is crucial to uncover the mechanisms driving LAM cell detachment, intravasation, and their impact on the lymphatic endothelial barrier. Understanding the interactions between LECs, fibroblasts, and LAM cells will provide critical insights into how LAM cells invade the lymphatic system and spread to other organs. This comprehensive understanding of LAM pathogenesis will be pivotal in developing innovative therapeutic strategies. In conclusion, our study reinforces the classification of LAM as a slow-growing neoplastic disease with a complex microenvironment driven by multiple cell types and signaling pathways. These findings not only open new avenues for therapeutic intervention but also underscore the need for continued research into the pathogenesis of LAM, with the ultimate goal of developing more effective treatments for this challenging disease.

## MATERIALS AND METHODS

Detailed methodology is found in the Online Data Supplement which can also be accessed at doi: 10.6084/m9.figshare.26880469

### Study design

The objective of this study was to create a co-culture 3-D model to evaluate cellular interactions between LAM associated fibroblasts and lymphatic endothelial cells in the pathogenesis of LAM. Our work included multiplexed immunohistochemistry and spatialomic analysis of LAM donor lung tissues, characterization of LAM-associated fibroblasts compared to healthy lung fibroblasts, quantitative evaluation of LAMF and HLF and co cultures with LEC in 3-D spheroid cultures and comparison of the effectiveness of current first line therapeutic, rapamycin, compared to a multikinase inhibitor on LAMF invasion. Cells and tissue samples were selected based on the diagnosis for the donor lung tissues, either having LAM or non-smokers with no prior history of chronic lung disease. Deidentified LAM lung samples were obtained from living donors at the time of lung transplantation through the National Disease Research Institute (NDRI, Philadelphia Pennsylvania) protocol number RKRV1. Informed consent was obtained by NDRI prior to acceptance of tissue donation for research. Identifying information were removed prior to use in accordance with institutional and NIH protocols (*48, 78*)Human lung tissue from subjects with no prior history of chronic lung disease was obtained International Institute for the Advancement of Medicine (IIAM), the CFF tissue procurement and cell culture core at UNC or the University of Iowa in collaboration with Dr. Kalpaj Parekh with approval from the Institutional Review Board (IRB) of the University of Southern California (USC) (Protocol number: #HS-18-00273). For histological experiments, 3 LAM lung tissues were used. For spatial omics analysis 2 LAM lung tissues were used. For all other experiments a minimum of 3 biological replicates from diseased (LAM) and controls (HLF) were used. For all 3-D culture analysis a minimum of 12 wells of spheroids were used per condition. Outliers were recorded and included.

### Invasion assay and image analysis

Fibroblast-LEC spheroids were created through the sequential addition of primary human pulmonary lymphatic microvascular endothelial cells maintained to the fibroblast spheroids. To monitor the growth and invasion daily phase contrast images were collected over 7 days. To evaluate morphology and phenotype of spheroids/organoids, an automated pipeline was developed using CellProfiler (available at https://cellprofiler.org). The pipeline was designed to enhance specific aspects of the phenotype and accurately identify both primary and secondary objects, including spheroid cores and spheroid sprouts (see **fig. S3**). The modified pipeline allowed for a more precise quantification of various features, including size and shape, and facilitated the analysis of the spheroid/organoid behavior under different experimental conditions. Images analysis parameters: 1) Compactness: A filled circle will have a compactness of 1, with irregular objects or objects with holes having a value greater than 1. 2) Perimeter: Total length of the perimeter of the objects image. 3) Solidity: Equals 1 for a solid object, or <1 for an object with holes or possessing a convex/irregular boundary. 3) Form factor: Calculated as 4*π*Area/Perimeter2. Equals 1 for a perfectly circular object, >1 for an irregular object.

### Visium 10X Spatial Transcriptomic Profiling

Spatial transcriptomic profiling was performed in collaboration with the Iowa NeuroBank Core in the Iowa Neuroscience Institute (INI) and the Genomics Division in the Iowa Institute of Human Genetics (IIHG), and followed manufacturer’s instructions (Doc. #CG000409, Rev. A). Briefly, two independent regions from one LAM lung donor were used to create 10 μM sections for quality control in the Comparative Pathology Laboratory (CPL) in the Department of Pathology, UIowa. Sections were H&E stained and regions of interest selected based on the presence of LAM nodules and Cysts. RNA quality was determined by extracting RNA with the Qiagen RNeasy FFPE kit (Qiagen, Hilden, Germany) and measuring RNA concentration using a Qubit fluorometer (ThermoFisher Scientific, Waltham, MA, USA) and quality determined by evaluating the samples using a 4200 Tapestation Agilent, Santa Clara, CA) in the Genomics Division of the Iowa Institute of Human Genetics. For sequencing, 10 μM FFPE sections were rehydrated, then adhered to the Visium Spatial Gene Expression Slide (PN-2000233, 10X Genomics) in a 42°C water bath. Samples were dried in a desiccation chamber and deparaffinized using Qiagen Deparaffinization Solution for 2 hours at 60°C. After H&E staining, brightfield tile scans of each complete section area were captured and stitched together using an Echo Revolution microscope with a 20X objective (software version 1.0.26.2). Decrosslinking was performed according to manufacturer’s protocol (Doc. #CG000407, Rev. D) and immediately hybridized to the Visium Human Transcriptome Probe Kit V1.0, which contained 19,144 genes targeted by 19,902 probes. Post-probe extension, sequencing library construction was performed using unique sample indices using the Dual Index Kit TS, Set A (PN-1000251) for Illumina-compatible sequencing. Paired-end sequencing (2×100) was performed an the Novaseq 4000.

### Statistical Analysis

All data is presented as mean ± S.E.M. For determining statistical significance among multiple groups in all experiments, a one-way analysis of variance (ANOVA) Kruskal-Wallis test was employed. Subsequently, post-hoc Dunn’s multiple comparisons test was utilized for pairwise comparisons between the groups. Statistical significance was considered at the 5% level, with a value of *p*<0.05 being considered statistically significant. Unless otherwise stated the data was collected from a minimum of 3 independent donor (N) and 3 independent experimental replicates (n).

## Supporting information

Supplementary Database S3

Supplementary Database S4

Supplementary Database S5

Supplementary Database S1

Supplemental Database S2

## List of Supplementary Materials

All supplemental materials and supporting raw data can be accessed through Figshare at doi:10.6084/m9.figshare.26880469.

Materials and Methods

Figures S1 to S6

Tables S1 to S4

Data files S1 to S5

## Acknowledgments

Mariah Leidinger, Comparative Pathology Core, University of Iowa prepared the tissue samples for spatial-transcriptomic processing. Visium data presented herein were obtained at the Iowa NeuroBank Core in the Iowa Neuroscience Institute (INI) and the Genomics Division in the Iowa Institute of Human Genetics (IIHG). Initial data processing and analysis was carried out by the Bioinformatics Division in the IIHG. INI is supported by the Roy J. Carver Charitable Trust and the University of Iowa Carver College of Medicine. IIHG is supported, in part, by the University of Iowa Carver College of Medicine. Opal Staining was performed in collaboration with the Translational Pathology Core Laboratory in the Department of Pathology and Laboratory Medicine at the UCLA David Geffen School of Medicine. The authors would like to acknowledge use of the University of Iowa Central Microscopy Research Facility (CMRF), a core resource supported by the University of Iowa Vice President for Research, and the Carver College of Medicine. *The use of the Zeiss LSM710 confocal microscope was supported by a Shared Instrumentation grant from the NIH under award number 1 S10 RR025439-01.* Finally, the authors would like to thank the Center for Gene Therapy Cell and Tissue Core, supported by NIH P30DK054759-22 and the CFF Tissue Procurement and Cell Culture Core at UNS, supported by CF BOUCE19R) and the NIH CFRTCC Cell Models Core at UNC supported by NIH P30DK065988cknowledgments follow the references and notes but are not numbered. Start with text that acknowledges non-author contributions and then complete each of the sections below as separate paragraphs.

## Funding

This study was supported by

The Hastings Foundation (ALR)

The LAM Foundation (ALR, LAM0154SG01-22-891077)

The German Research Foundation (SKG, KO5803/2-1)

NIH/NHLBI (VPK, RO1 HL141462, RO1 HL158737, RO1151647, and U01HL131022)

## Author contributions

Conceptualization: SKG, ALR

Methodology: SKG K, EL, LKG, ALR

Investigation: SKG, EL, LKG, BAC, SM, NAC, JCN, ALR

Visualization: SKG, EL, LKG, SM, JCN, ALR

Funding acquisition: SKG, VPK, ALR

Project administration: SKG, ALR

Supervision: SKG, BZ, ALR

Writing – original draft: SKG, ALR

Writing – review & editing: SKG, EL, LKG, VPK, ALR

## Data and materials availability

All data associated with this study are present in the paper or the Supplementary Materials (Doi: 10.6084/m9.figshare.26880469). The data spatial-transcriptomics datasets, including raw sequencing data and processed files, have been deposited at GEO under the series reference GSE234885 (datasets GSM7476184 and GSM7476185). This paper includes original coding which is available in the Git-Hub repository (https://github.com/gautam-lk/RyanLab_LAM). In addition, all raw data is available at 10.6084/m9.figshare.23464976. Any additional information required to reanalyze the data reported in this paper is available from the lead contact upon request.

