## Supplemental Data File for "Targeting Fibroblast-Endothelial Interactions in LAM Pathogenesis: 3D Spheroid and Spatial Transcriptomic Insights for Therapeutic Innovation"

**Affiliations:**
[1]Hastings Center for Pulmonary Research, Division of Pulmonary, Critical Care and Sleep Medicine, Department of Medicine, University of Southern California; Los Angeles, CA, USA
[2]Department of Stem Cell Biology and Regenerative Medicine, University of Southern California; Los Angeles, CA, USA
[3]Department of Internal Medicine II, Infectious Diseases, University Hospital Frankfurt; Goethe University Frankfurt, Frankfurt am Main, Germany
[4]Department of Anatomy and Cell Biology, Carver College of Medicine, University of Iowa; Iowa City, IA, USA
[5]Helmholtz Pioneer Campus and Institute of Biological and Medical Imaging; Helmholtz Zentrum München, Neuherberg, Germany
[6]Division of Pulmonary and Critical Care Medicine, Lung Biology Institute, Perelman School of Medicine, University of Pennsylvania; Philadelphia, PA, USA

**One Sentence Summary:** Using 3D spheroids and spatial transcriptomics, we identified LAMFs and LECs as key contributors to LAM, with bFGF and VEGF-A as potential therapeutic targets

**Abstract:** Lymphangioleiomyomatosis (LAM) is a progressive lung disease with limited treatments, largely due to an incomplete understanding of its pathogenesis. Lymphatic endothelial cells (LECs) invade LAM cell clusters, which include HMB-45-positive epithelioid cells and smooth muscle α-actin-expressing LAM-associated fibroblasts (LAMFs). Recent evidence shows that LAMFs resemble cancer-associated fibroblasts, with LAMF-LEC interactions contributing to disease progression. To explore these mechanisms, we used spatial transcriptomics on LAM lung tissues and identified a gene cluster enriched in kinase signaling pathways linked to myofibroblasts and co-expressed with LEC markers. Kinase arrays revealed elevated PDGFR and FGFR in LAMFs. Using a 3D co-culture spheroid model of primary LAMFs and LECs, we observed increased invasion in LAMF-LEC spheroids compared to non-LAM fibroblasts. Treatment with Sorafenib, a multikinase inhibitor, significantly reduced invasion, outperforming Rapamycin. We also confirmed TSC2-null AML cells as key VEGF-A secretors, which was suppressed by Sorafenib in both AML cells and LAMFs. These findings highlight VEGF-A and bFGF as potential therapeutic targets and suggest multikinase inhibition as a promising strategy for LAM.

### Supplemental Materials and Methods

**Human lung tissue and primary cell culture**:_Primary human cells were derived from donated explanted lungs as previously described *(1, 2)*. De-identified lung tissue samples from patients with advanced LAM disease who had undergone lung transplantation were received from the National Disease Research Interchange (NDRI). Use of these tissues does not constitute human subjects research since all donor tissue is harvested anonymously and de-identified. LAM-cells were grown in FBM-2 media (Lonza Walkersville, Inc.) supplemented with hFGF2 (human recombinant fibroblast growth factor), Insulin, FBS (fetal bovine serum), gentamicin as per the manufacturers instruction and maintained at 37$^o$C in a humidified atmosphere with 5% $CO_2$. Human lung tissue was obtained from subjects with no prior history of chronic lung disease for fibroblast isolation. A minimum of three independent donor cells were used for LAM-fibroblasts (LAMF) and Human Lung Fibroblasts (HLF), donor information is detailed in **Supplemental Table S1**.

**Spheroid culture*:*** Spheroid Formation: Spheroids were generated from primary human bronchial SMC maintained in VascuLife Media (hBrSMC, Lifeline Cell Technology, San Diego, CA), LAMF or HLF. Cells were re-suspended using TrypLE (Thermo Fisher Scientific) for 5 minutes at 37°C, 5% $CO_2$ and collected by centrifugation at 1300 rpm for 5 minutes and 500K cells distributed into each well of an AggreWell™800 plate (24 well) (Stem Cell Technologies) as recommended by the manufacturer. Multicellular Spheroid Formation: Fibroblast-LEC spheroids were created through the addition of primary human pulmonary lymphatic microvascular endothelial cells maintained in Endothelial Growth Medium (LECs, Angio-Proteomie, Atlanta, GA) to the fibroblast spheroids. A sequential culture approach was performed to create a LAM-nodule comprising of a core of LAM-cells cultured with LECs. On day 1 harvested HLF and LAMF were resuspended in warm 15 µM CellTracker$^{TM}$-CMFDA (green) and incubated for 30 min. Cells were then centrifuged and resuspended in culture media. Cells were then seeded on AggreWell™800 plates and centrifuged to form spheroids after 24h. On day 2 LECs were lifted and labelled with CellTracker™ Red CMTPX. Labelled LECs were then seeded on the pre-formed spheroids and centrifuged at 100 x g for 3 min.

**IF OPAL staining**: Opal Multiplex Staining was used to visualize multiple proteins in a single tissue sample with same species antibodies as previously described *(3, 4)*. Samples were either stained manually or automatically on Leica Bond RX (BOND-MAX; Leica Biosystems) following manufacturers protocols. For both methods the preparation and staining steps used the same steps. Briefly, tissue slides were prepared by deparaffinizing, dewaxing and dehydrating through ethanol incubation and rehydration with PBS washing. Epitope retrieval was performed using a microwave and AR6/AR9 buffer manually or BOND Antigen Retrieval buffer on the automated system. After blocking, primary antibodies were incubated overnight at 4°C, followed by secondary antibody incubation at RT with HRP (RNASCOPE Multiplex Fl V2 HRP Blocker, 323107) labeled antibodies. Opal 520 or Opal 570 was incubated for 10 minutes at RT for staining. For additional antibodies the sample was treated with HRP blocker and incubated in oven at 40°C for 40 minutes and the staining steps were repeated. Finally, the samples were washed with TBST, and cover slipped with a DAPI mounting medium. The antibodies used in this study are listed in **Supplemental Table S2**. Images were visualized and evaluated using Phenochart (Akoya Biosciences, Menlo Park, CA)

**Invasion assay and image analysis:** Spheroids were generated in AggreWell™ dishes as per manufacturer protocol and embedded in Matrigel as follows. The bottom of each well was coated in 20 µL of a 50/50 mixture of Matrigel and high-glucose DMEM (Gibco, #11965092). Spheroids were re-suspended in 50/50 Matrigel/DMEM and placed in each well. 50 µL of 50/50 Matrigel/DMEM was used to cover the spheroids. The Matrigel was allowed to set at 37C for 5 minutes between each step. Fibroblast medium (10% Fetal Bovine Serum (Gibco, # 26140079), 1% GlutaMax (Gibco, #35050061), and 1% Penicillin-Streptomycin (Gibco, #15140122) in high-glucose DMEM) was added to the apical side of the gel. Fibroblast-LEC spheroids were created through the sequential addition of primary human pulmonary lymphatic microvascular endothelial cells (Angio-Proteomie, #cAP-0039) to the fibroblast spheroids and maintained in 50/50 fibroblast medium and Endothelial Growth Medium (Angio-Proteomie, #cAP-02). To monitor the growth and invasion daily phase contrast images were collected over 72 hours. To evaluate morphology and phenotype of spheroids/organoids, an automated pipeline was developed using CellProfiler (available at https://cellprofiler.org). The pipeline was designed to enhance specific aspects of the phenotype and accurately identify both primary and secondary objects, including spheroid cores and spheroid sprouts (see **Supplemental Figure S3**). The modified pipeline allowed for a more precise quantification of various features, including size and shape, and facilitated the analysis of the spheroid/organoid behavior under different experimental conditions. Images analysis parameters: 1) **Compactness:** A filled circle will have a compactness of 1, with irregular objects or objects with holes having a value greater than 1. 2) **Perimeter:** Total length of the perimeter of the objects image. 3) **Solidity:** Equals 1 for a solid object, or <1 for an object with holes or possessing a convex/irregular boundary. 3) **Form factor:** Calculated as 4*π*Area/Perimeter2. Equals 1 for a perfectly circular object , >1 for an irregular object and phenotype of spheroids/organoids, an automated pipeline was developed using CellProfiler (available at https://cellprofiler.org). The pipeline was designed to enhance specific aspects of the phenotype and accurately identify both primary and secondary objects, including spheroid cores and spheroid sprouts (see **Supplemental Figure S3**). The modified pipeline allowed for a more precise quantification of various features, including size and shape, and facilitated the analysis of the spheroid/organoid behavior under different experimental conditions. Images analysis parameters: 1) **Compactness:** A filled circle will have a compactness of 1, with irregular objects or objects with holes having a value greater than 1. 2) **Perimeter:** Total length of the perimeter of the objects image. 3) **Solidity:** Equals 1 for a solid object, or <1 for an object with holes or possessing a convex/irregular boundary. 3) **Form factor:** Calculated as 4*π*Area/Perimeter2. Equals 1 for a perfectly circular object , >1 for an irregular object.

**Immunocytochemistry:** Adherent 2D samples were fixed in PFA 0.4% (paraformaldehyde) for 15 minutes at RT. Cells were permeabilized with Triton X-100 0.5% for 5 min at RT and blocked in buffer containing 5% normal donkey serum and 2% BSA for a minimum of 1 hour at RT. Incubation in primary antibody was performed overnight at 4°C. Secondary antibody was applied for 2 hours at RT in the dark. Finally, samples were mounted using ProLong(TM) Diamond Antifade Mountant with DAPI (Invitrogen; #P36962). 3D spheroid samples were fixed in PFA 0.4% (paraformaldehyde) for 15 minutes at RT. Spheroids were permeabilized and blocked with a buffer containing 0.3% Triton X-100 and 5% BSA in PBS overnight at 4°C. Secondary antibody was applied overnight at 4°C. Finally, samples were mounted using ProLong(TM) Diamond Antifade Mountant (Invitrogen; #P36961). The antibodies used for staining are included in Supplemental Table S2. Cells and spheroids were imaged in the University of Iowa Anatomy and Cell Biology Department, on the Zeiss LSM980 confocal microscope and data was analyzed in Fiji 2.13.1 or using Imaris 10 (Oxford Instruments software)

**Western Blotting/Immunoblotting:** Cells were grown to confluency and lysed with RIPA Lysis Buffer (Sigma-Aldrich, #R0278) with protease inhibitor (Roche, #11836170001) and phosphatase inhibitor (Roche, #04906845001). Protein content was measured using a Micro BCA Protein Assay Kit (Thermo Scientific, #23235) according to the manufacturer's protocol. Samples were prepared using 10 ug of total protein in LDS Sample Buffer (Invitrogen, #NP0007). SDS-PAGE was run using a 4-12% Bis-Tris Gel (Invitrogen, #NW04120), and transferred to a 0.2 um PVDF membrane (Bio-Rad, #1620177) using an Invitrogen Mini Gel Tank (Invitrogen, #A25977) as per the manufacturer's' protocol. Membranes were blocked using Blocking Buffer (5% Non-fat dry milk, 0.1% Tween 20, in 1X TBS). Incubation in primary antibodies was performed overnight at 4°C. Incubation with secondary antibodies was performed for 2 hours at RT. Finally, membranes were visualized using the ECL(TM) Prime Western Blotting Detection Reagents (Cytiva, #RPN2232) and imaged using an Amersham Imager 600 series imager. For repeated blots, membranes were stripped using Restore(TM) PLUS Western Blot Stripping Buffer (Thermo Scientific, #46340) and re-blocked with Blocking Buffer before re-use. The antibodies used for staining are included in **Supplemental Table S2**.D*ata was analyzed in Fiji 2.13.1.*

**RNA isolation and qPCR:** Cells were grown to confluency and RNA was isolated using the Direct-zol RNA Miniprep Kit (Zymo Research, #R2052) according to manufacturer's protocol. Total RNA content was quantified using the NanoDrop™ One/OneC Microvolume UV-Vis Spectrophotometer (Thermo Scientific, #ND-ONE-W). cDNA was generated from 1000 ng of RNA using the High-Capacity cDNA Reverse Transcription Kit (Applied Biosystems, #4368814) using the recommended protocol scaled to the desired volume for each experiment. PowerUp™ SYBR™ Green Master Mix for qPCR (Applied Biosystems, #A25742) was used for amplification and primers were added to a final concentration of 1 µM, and the qPCR reactions were carried out in a 364-well plate in a 10 µL reaction volume. The primers used are included in **Supplemental Table S3**. qPCR was run on the QuantStudio 5 Real-Time PCR System (Applied Biosystems,

#A28140). Samples were held at 50°C for 2 min then denatured at 95°C for 10 min. Samples were held at 95°C for 15 sec, then 60°C for 1 min, and this step was repeated 40 times. Finally, the samples were brought to 95°C for 15 sec, 60°C for 1 min, and 95°C for 15 sec. Data was collected at each 60°C step. Data was normalized to the housekeeping gene (RPLPO) and expressed as a fold change relative to the average of the control samples.

**Kinase array:** The kinase array and primary analysis was performed by Full moon-Biosystems (Sunnyvale, CA) following the manufacturers protocol and detailed in the Online Supplemental Methods. Briefly, after the protein concentration in the samples was measured 100 µL of DMF (Dimethylformamide) was added to 1 mg of Biotin Reagent to give a final concentration of 10 µg/µL. The samples were mixed and incubated at room temperature (RT) for two hours in a total of 75 µl of buffer and 3 µl of Biotin/DMF mix. 35 µL of Stop Reagent was added and incubated for an additional 30 minutes at RT. For the coupling process the antibody microarray was first blocked at RT for 40 minutes, rinsed in Milli-Q water and incubated in 6mL coupling solution on an orbital shaker for 2 hours at RT. After washing, the slide was incubated Cy3-Streptavidin solution an orbital shaker for 45 minutes at RT in the dark. After another wash the slide was dried with compressed nitrogen and scanned on Axon GenePix Array Scanner.

**Visium 10X Spatial Transcriptomic Profiling:** Spatial transcriptomic profiling was performed in collaboration with the Iowa NeuroBank Core in the Iowa Neuroscience Institute (INI) and the Genomics Division in the Iowa Institute of Human Genetics (IIHG), and followed manufacturer's instructions (Doc. #CG000409, Rev. A). Briefly, two independent regions from one LAM lung donor were used to create 10 µM sections for quality control in the Comparative Pathology Laboratory (CPL) in the Department of Pathology, UIowa. Sections were H&E stained and regions of interest selected based on the presence of LAM nodules and Cysts. RNA quality was determined by extracting RNA with the Qiagen RNeasy FFPE kit (Qiagen, Hilden, Germany) and measuring RNA concentration using a Qubit fluorometer (ThermoFisher Scientific, Waltham, MA, USA) and quality determined by evaluating the samples using a 4200 Tapestation Agilent, Santa Clara, CA) in the Genomics Division of the Iowa Institute of Human Genetics. For sequencing, 10 µM FFPE sections were rehydrated, then adhered to the Visium Spatial Gene Expression Slide (PN-2000233, 10X Genomics) in a 42ºC water bath. Samples were dried in a desiccation chamber and deparaffinized using Qiagen Deparaffinization Solution for 2 hours at 60ºC. After H&E staining, brightfield tile scans of each complete section area were captured and stitched together using an Echo Revolution microscope with a 20X objective (software version 1.0.26.2). Decrosslinking was performed according to manufacturer's protocol (Doc. #CG000407, Rev. D) and immediately hybridized to the Visium Human Transcriptome Probe Kit V1.0, which contained 19,144 genes targeted by 19,902 probes. Post-probe extension, sequencing library construction was performed using unique sample indices using the Dual Index Kit TS, Set A (PN-1000251) for Illumina-compatible sequencing. Paired-end sequencing (2x100) was performed an the Novaseq 4000.

**Bioinformatic Analysis of Spatial-transcriptomics:** Initial data analysis was carried out in the Bioinformatics Division of the Iowa Institute of Human Genetics (IIHG). Briefly, Illumina BCL data was demultiplexed and converted to FASTQ format with Illumina's BCL2Fastq (v2.20) on IIHG servers. The resulting FASTQ sequencing data was then processed using the SpaceRanger "count" pipeline, v2.0.1, (10X Genomics) on the University of Iowa's Argon high-performance computing cluster. SpaceRanger was run with 32 local cores and 128GB of RAM on a single 56-core standard-memory node. SpaceRanger "count" carries out read alignment against the GRCh38 reference genome and the probe set which was specified as "Visium_Human_Transcriptome_Probe_Set_v1.0_GRCh38-2020-A.csv" (available as a download from 10X Genomics) and generates "feature-spot" matrices for downstream analysis. The pipeline also performs automatic tissue detection and fiducial alignment from the images." Primary data was evaluated, and analysis was performed using Loupe Browser v6.1 from .cloupe files created by the SpaceRanger pipeline, v2.0. Further, a list of significant DEGs from all the clusters was generated from filtered matrix (H5) files using the Seurat 10X Visium analysis pipeline. SeuratObject from both datasets (LAM_D1 and LAM_D2) were integrated *(5, 6)* and re-clustered for enriched and DEGs (with Log2 fold change ≥ 1.5) from LAM-core cluster was exported for mapping to LungMAP. The average expression values of all significant genes and a subset of 50 genes in the LAM-core were plotted across all other clusters as heatmaps. The gene list was further refined to plot a heatmap against the Fullmoon kinase assay data using Euclidean clustering method. Heatmaps were prepared using the pHeatmap R-package *(7)* in RStudio (RStudio Team, 2020; R Core Team, 2021).

LAM-core significant DEG were also mapped for their distribution across the sorted cell type annotations based on human single cells dataset in the Lung Gene Expression Analysis (LEGA) Web portal *(8, 9)*. Code and processed data used for spatial mapping and Heatmap is available in the Git-Hub repository (https://github.com/gautam-lk/RyanLab_LAM). The cellranger matrix output file was used to predict cell type with azimuth application *(10)* using Human Lung Single cell dataset Lung v2 HLCA *(11)*. LAM-core specific marker genes were used as query features to identify the LAM cluster cell type. A reference cell list was generated based on Azimuth cell type prediction for mapping of features and generating cell specific spatial image. A reference-based probabilistic model was used for mapping the UMIs on the tissue sections for cell type deconvolution analysis *(12)*. The single cell and single nuclei data were processed for Seurat integrative analysis *(13)*. The LAM-Core signature genes (GEO datasets: GSM4035465, GSM4035466, GSM4035467 and GSM4035468) were used for identification of LAM core cluster genes *(14)*.

**Ingenuity Pathway Analysis (IPA)**: Analysis of the LAM-core spatialomics expression clusters from the integrated datasets was completed using IPA (QIAGEN Inc., https://www.qiagenbioinformatics.com/products/ingenuity-pathway-analysis). Data with ≥1.5 $Log_2$-fold change was filtered to perform core analysis using "expression analysis" criteria. The gene population for calculating the p-value was limited only to the Qiagen Ingenuity Knowledge Base, with both bi-directional relationships among genes impacting Canonical pathways, interaction networks and upstream regulators. For Canonical pathways and casual network analysis to identify master regulators related to any disease, functions genes or chemical selections, "Kinase" was chosen only for comparison with the Full moon Kinase Assay dataset.

**Statistical Analysis:** All data is presented as mean ± S.E.M. For determining statistical significance among multiple groups in all experiments, a one-way analysis of variance (ANOVA) Kruskal-Wallis test was employed. Subsequently, post-hoc Dunn's multiple comparisons test was utilized for pairwise comparisons between the groups. Statistical significance was considered at the 5% level, with a value of $p<0.05$ being considered statistically significant. Unless otherwise stated the data was collected from a minimum of 3 independent donor (N) and 3 independent experimental replicates (n).

**Study Approval:** Deidentified LAM lung samples were obtained from living donors at the time of lung transplantation through the National Disease Research Institute (NDRI, Philadelphia Pennsylvania) protocol number RKRV1. Informed consent was obtained by NDRI prior to acceptance of tissue donation for research. Identifying information were removed prior to use in accordance with institutional and NIH protocols (43, 51).Human lung tissue from subjects with no prior history of chronic lung disease was obtained International Institute for the Advancement of Medicine (IIAM), the CFF tissue procurement and cell culture core at UNC or the University of Iowa in collaboration with Dr. Kalpaj Parekh with approval from the Institutional Review Board (IRB) of the University of Southern California (USC) (Protocol number: #HS-18-00273).

**SUPPLEMENTAL TABLES**

**Supplemental Table S1: Human Lung Tissue Donor Information**

| Donor Code | Laboratory | Cell Description | Age | Sex | Ethnicity | Other information |
|---|---|---|---|---|---|---|
| LAMF_A | Krymskaya | LAM associated fibroblasts | 57 | F | Cauc. | LAM dx by CAT scan<br>PMH: Arthritis, heart disease/ irregular heartbeat.<br>No tobacco or illicit drug use. |
| LAMF_B | Krymskaya | LAM associated fibroblasts<br><br>LAM Tissue Blocks | 70 | F | Cauc. | LAM dx by open lung biopsy<br>PMH: high blood pressure, acid reflux, COPD, high cholesterol, anxiety, pneumonia, Hashimoto's Thyroiditis.<br>Smoked cigarettes 2 packs/day for 33 years.<br>Not a current smoker.<br>Surgery history: hysterectomy |
| LAMF_C | Krymskaya | LAM associated fibroblasts | 67 | F | Cauc. | LAM dx unknown method<br>PMH: high blood pressure, high cholesterol (mildly), depression, kidney trouble |
| LAM_D | Krymskaya | LAM Tissue blocks | 50 | F | Cauc./ Black | LAM dx unknown method<br>PMH: Asthma, Arthritis, COPD d/t LAM, Anxiety/ Depression, Hx of pneumonia |
| LAM_E | Krymskaya | LAM Tissue blocks | 45 | F | Cauc. | LAM dx by CT scan.<br>PMH: Asthma, pneumonia<br>Non-smoker. |
| HLF_1 | Ryan | HLF | U/K | U/K | U/K | Non-smoker, no chronic lung disease |
| HLF_2 | Ryan | HLF | U/K | U/K | U/K | Non-smoker, no chronic lung disease |
| HLF_3 | Ryan | HLF | U/K | U/K | U/K | Non-smoker, no chronic lung disease |
| HLF_4 | Ryan | HLF | 19 | M | Hisp. | Non-smoker, no chronic lung disease |
| HLF_5 | Ryan | HLF | 24 | F | Cauc. | Non-smoker, no chronic lung disease |

Cauc., Caucasian; COPD, chronic obstructive pulmonary disease; d/t, due to; dx, diagnosis; F, female; Hisp., Hispanic; Hx, history; HLF, human lung fibroblasts; LAMF, LAM fibroblasts; PMH, past medical history; U/K, unknown

**Supplemental Table S2: Detailed information for all antibodies used in this study.**

| Antibodies | Company/Cat.# | Host | Application/Dilution |
|---|---|---|---|
| Primary Antibodies | | | |
| PDGFRa | Cell Signaling/#3174S | Rabbit | WB; 1:1000 |
| αSMA | Thermo Fisher/#MA5011547 | Mouse | WB; 1:500, IF; 1:200 |
| HMB-45 | Agilent DAKO/#GA052 | Mouse | 1:50 |
| VEGFR3 | R&D/#AF349 | Goat | 1:100 |
| D2-40 | Biocare Medical/#CM266A | Mouse | 1:100 |
| PROX1 | R&D/#AF2727 | Goat | 1:50 |
| CD144 | Invitrogen/#14-1449-82 | Mouse | 1:100 |
| LYVE-1 | Invitrogen/#MA5-32512 | Rabbit | 1:50 |
| PDGFRβ | Cell Signaling/#3169 | Rabbit | WB; 1:1000, IF; 1:200 |
| CAMKIID | Invitrogen/#PA5-22168 | Rabbit | WB; 1:100 |
| Hic-5 (TGFB1I1) | SantaCruz; #SC-271353 | Mouse | WB; 1:1000 |
| B-actin | GeneTex; GTX109639 | Rabbit | WB; 1:2000 |
| mTOR | Cell Signaling; #2972 | Rabbit | WB; 1:500 |
| Phospho-mTOR | Cell Signaling; #5526 | Rabbit | WB; 1:1000 |
| P70 S6K | Cell Signaling; #34475s | Rabbit | WB; 1:1000 |
| Phospho-P70 S6K | Cell Signaling; #9234 | Rabbit | WB; 1:1000 |
| Secondary Antibodies | | | |
| Anti-mouse-AF647 | Invitrogen/#A31571 | Donkey | 1:500 |
| Anti-goat-AF488 | Invitrogen/#A11055 | Donkey | 1:500 |
| Anti-rabbit-AF568 | Invitrogen/#A10042 | Donkey | 1:500 |
| Anti-rabbit-AF594 | Invitrogen/#A11036 | Donkey | 1:500 |
| OPAL 520 | Akoya Biosciences/#NEL810001KT | | As per manuracturer's protocol |
| OPAL 570 | | | |
| OPAL 620 | | | |
| OPAL 690 | | | |
| Goat Anti-Rabbit IgG HRP | Abcam; #ab6721 | Goat | WB; 1:1000 |
| Horse anti-Mouse HRP | Cell Signaling; #7076s | Horse | WB; 1:1000 |

**Supplemental Table S3: Primers used in this study.**

| Gene | Primer F' | Primer R' | Amplicon Size |
|---|---|---|---|
| PDGFRβ | CAAGGACACCATGCGGCTTC | AGCAGGTCAGAACGAAGGTG | 173 |
| Hic-5 (TGFB1I1) | TCTCCTCTTCCAGCGGTGTCTTT | TTCTGCTCTCCTGAAGCCACCTT | 142 |
| RPLPO | CCGTGATGCCCAGGGAAGAG | GCATCTGCTTGGAGCCCACA | 126 |
| FAP | GTCCTGGCTTCAGCTTCCAA | AGGGCGTAAGACAATGCACA | 202 |
| TGFB1 | TGGTGGAAACCCACAACGAA | GAGCAACACGGGTTCAGGTA | 113 |
| ACTA2 | CTTTGGCTTGGCTTGTCAGG | GTTGGAGAGAGGAGCGGAAC | 52 |

**Supplemental Table S4: Azimuth App Lung V2 (Cell type annotation markers).**

| Date Assessed: 22 July 2024 | |
|---|---|
| **Annotation level 3** | |
| **Label** | **Markers** |
| Macrophages | C1QB, C1QA, HLA-DRB1, AIF1, LYZ, CTSZ, CTSL, FCER1G, C1QC, LAPTM5 |
| Innate lymphoid cell NK | GZMA, CD7, CCL4, CST7, NKG7, GNLY, CTSW, CCL5, GZMB, PRF1 |
| AT2 | NAPSA, SFTPD, PEBP4, SLC39A8, SLC34A2, CYB5A, MUC1, S100A14, SFTA2, SFTA3 |
| Basal | KRT15, SFN, IGFBP2, SERPINF1, ADIRF, MT1X, BCAM, KRT5, KRT17, S100A2 |
| EC venous | VWF, MGP, RAMP3, GNG11, RAMP2, SPARCL1, IGFBP7, CLEC14A, ACKR1, PECAM1 |
| T cell lineage | CD69, CORO1A, CXCR4, CD3E, TRBC2, TRAC, CD2, CCL5, IL32, CD3D |

| | |
|---|---|
| EC arterial | SPARCL1, SOX17, IFI27, TM4SF1, A2M, CLEC14A, GIMAP7, CRIP2, CLDN5, PECAM1 |
| Fibroblasts | BGN, DCN, MGP, SPARC, CALD1, LUM, COL6A1, IGFBP7, COL1A2, C1S |
| AT1 | SFTA2, CEACAM6, FXYD3, CAV1, TSPAN13, KRT7, ADIRF, HOPX, AGER, EMP2 |
| Multiciliated lineage | SNTN, FAM229B, TMEM231, C5orf49, C12orf75, GSTA1, C11orf97, RP11-356K23.1, CD24, RP11-295M3.4 |
| B cell lineage | TSC22D3, ISG20, IGHA1, CORO1A, IGKC, IGLC2, LTB, CD79A, CD37, MS4A1 |
| Secretory | KRT19, CEACAM6, WFDC2, TACSTD2, CXCL17, BPIFB1, MUC1, CP, KRT7, PIGR |
| SM activated stress response | C11orf96, HES4, PLAC9, FLNA, KANK2, TPM2, PLN, SELM, GPX3, LBH |
| Monocytes | FCER1G, S100A9, S100A12, VCAN, COTL1, S100A8, CD14, LST1, FCN1, AIF1 |
| Submucosal Secretory | CCL28, PIP, ZG16B, PIGR, SEC11C, MARCKSL1, SELM, LTF, TCN1, AZGP1 |
| EC capillary | IFI27, TM4SF1, CLEC14A, CDH5, PECAM1, HYAL2, SPARCL1, EGFL7, CLDN5, AQP1 |
| Lymphatic EC mature | PPFIBP1, GNG11, RAMP2, CCL21, MMRN1, IGFBP7, SDPR, TM4SF1, CLDN5, ECSCR |
| Rare | FOXI1, HEPACAM2, ATP6V1A, EPCAM, KRT7, TMEM61, ASCL3, CLCNKB, SEC11C, CD24 |
| Lymphatic EC differentiating | AKAP12, TFF3, SDPR, CLDN5, TCF4, TFPI, TIMP3, GNG11, CCL21, IGFBP7 |
| Dendritic cells | CORO1A, MS4A6A, ITGB2, GPR183, HLA-DRB1, HLA-DPB1, HLA-DPA1, HLA-DQB1, HLA-DQA1, HLA-DMA |
| Myofibroblasts | CALD1, CYR61, TAGLN, MT1X, PRELP, TPM2, GPX3, CTGF, IGFBP5, SPARCL1 |
| Mast cells | VWA5A, RGS13, C1orf186, HPGDS, CPA3, GATA2, MS4A2, KIT, TPSAB1, TPSB2 |
| Fibromyocytes | NEXN, ACTG2, LMOD1, IGFBP7, PPP1R14A, DES, FLNA, TPM2, PLN, SELM |
| Lymphatic EC proliferating | S100A16, TUBB, HMGN2, COX20, LSM2, HMGN1, ARPC1A, ECSCR, EID1, MARCKS |
| **Annotation level: Finest** | |
| **Label** | **Markers** |
| Alveolar macrophages | MS4A7, C1QA, HLA-DQB1, HLA-DMA, HLA-DPB1, HLA-DPA1, ACP5, C1QC, CTSS, HLA-DQA1 |
| NK cells | GZMA, CD7, CCL4, CST7, NKG7, GNLY, CTSW, CCL5, GZMB, PRF1 |
| AT2 | SEPP1, PGC, NAPSA, SFTPD, SLC34A2, CYB5A, MUC1, S100A14, SFTA2, SFTA3 |
| Alveolar Mφ CCL3+ | MCEMP1, UPP1, HLA-DQA1, C5AR1, HLA-DMA, AIF1, LST1, LINC01272, MRC1, CCL18 |
| Suprabasal | PRDX2, KRT19, SFN, TACSTD2, KRT5, LDHB, KRT17, KLK11, S100A2, SERPINB4 |
| Basal resting | CYR61, PERP, IGFBP2, KRT19, KRT5, KRT17, KRT15, S100A2, LAMB3, BCAM |
| EC venous pulmonary | VWF, MGP, GNG11, RAMP2, SPARCL1, IGFBP7, IFI27, CLDN5, ACKR1, AQP1 |
| CD8 T cells | CD8A, CD3E, CCL4, CD2, CXCR4, GZMA, NKG7, IL32, CD3D, CCL5 |
| EC arterial | SPARCL1, SOX17, IFI27, TM4SF1, A2M, CLEC14A, GIMAP7, CRIP2, CLDN5, PECAM1 |
| Peribronchial fibroblasts | IGFBP7, COL1A2, COL3A1, A2M, BGN, DCN, MGP, LUM, MFAP4, C1S |
| CD4 T cells | CORO1A, KLRB1, CD3E, LTB, CXCR4, IL7R, TRAC, IL32, CD2, CD3D |
| AT1 | SFTA2, CEACAM6, FXYD3, CAV1, TSPAN13, KRT7, ADIRF, HOPX, AGER, EMP2 |
| Multiciliated (non-nasal) | SNTN, FAM229B, TMEM231, C5orf49, C12orf75, GSTA1, C11orf97, RP11-356K23.1, CD24, RP11-295M3.4 |
| Plasma cells | ITM2C, TNFRSF17, FKBP11, IGKC, IGHA1, IGHG1, CD79A, JCHAIN, MZB1, ISG20 |
| Goblet (nasal) | KRT7, MUC1, MUC5AC, MSMB, CP, LMO7, LCN2, CEACAM6, BPIFB1, PIGR |
| Club (nasal) | ELF3, C19orf33, KRT8, KRT19, TACSTD2, MUC1, S100A14, CXCL17, PSCA, FAM3D |
| SM activated stress response | C11orf96, HES4, PLAC9, FLNA, KANK2, TPM2, PLN, SELM, GPX3, LBH |
| Classical monocytes | LST1, IL1B, LYZ, COTL1, S100A9, VCAN, S100A8, S100A12, AIF1, FCN1 |
| Monocyte derived Mφ | LYZ, ACP5, TYROBP, LGALS1, CD68, AIF1, CTSL, EMP3, FCER1G, LAPTM5 |

| | |
|---|---|
| Alveolar Mφ proliferating | H2AFV, STMN1, LSM4, GYPC, PTTG1, KIAA0101, FABP4, CKS1B, UBE2C, HMGN2 |
| Club (non-nasal) | SCGB3A1, CYP2F1, GSTA1, HES4, TSPAN8, TFF3, MSMB, BPIFB1, SCGB1A1, PIGR |
| SMG serous (bronchial) | AZGP1, ZG16B, PIGR, NDRG2, LPO, C6orf58, DMBT1, PRB3, FAM3D, RP11-1143G9.4 |
| EC venous systemic | VWF, MGP, GNG11, PLVAP, RAMP2, SPARCL1, IGFBP7, A2M, CLEC14A, ACKR1 |
| Non classical monocytes | PSAP, FCGR3A, FCN1, CORO1A, COTL1, FCER1G, LAPTM5, CTSS, AIF1, LST1 |
| EC general capillary | EPAS1, GNG11, IFI27, TM4SF1, EGFL7, AQP1, VWF, FCN3, SPARCL1, CLDN5 |
| Adventitial fibroblasts | COL6A2, SFRP2, IGFBP7, IGFBP6, COL3A1, C1S, MMP2, MGP, SPARC, COL1A2 |
| Lymphatic EC mature | PPFIBP1, GNG11, RAMP2, CCL21, MMRN1, IGFBP7, SDPR, TM4SF1, CLDN5, ECSCR |
| EC aerocyte capillary | EMCN, HPGD, IFI27, CA4, EGFL7, AQP1, IL1RL1, SPARCL1, SDPR, CLDN5 |
| Smooth muscle | PRKCDBP, NDUFA4L2, MYL9, ACTA2, MGP, CALD1, TPM1, TAGLN, IGFBP7, TPM2 |
| Alveolar fibroblasts | LUM, COL6A1, CYR61, C1R, COL1A2, MFAP4, A2M, C1S, ADH1B, GPX3 |
| Multiciliated (nasal) | RP11-356K23.1, EFHC1, CAPS, ROPN1L, RSPH1, C9orf116, TMEM190, DNALI1, PIFO, ODF3B |
| Goblet (bronchial) | MUC5AC, MSMB, PI3, MDK, ANKRD36C, TFF3, PIGR, SAA1, CP, BPIFB1 |
| Neuroendocrine | UCHL1, TFF3, APOA1BP, CLDN3, SEC11C, NGFRAP1, SCG5, HIGD1A, PHGR1, CD24 |
| Lymphatic EC differentiating | AKAP12, TFF3, SDPR, CLDN5, TCF4, TFPI, TIMP3, GNG11, CCL21, IGFBP7 |
| DC2 | ITGB2, LAPTM5, HLA-DRB1, HLA-DPB1, HLA-DPA1, HLA-DMB, HLA-DQB1, HLA-DQA1, HLA-DMA, LST1 |
| Transitional Club AT2 | CXCL17, C16orf89, RNASE1, KRT7, SCGB1A1, PIGR, SCGB3A2, KLK11, SFTA1P, FOLR1 |
| DC1 | HLA-DPA1, CPNE3, CORO1A, CPVL, C1orf54, WDFY4, LSP1, HLA-DQB1, HLA-DQA1, HLA-DMA |
| Myofibroblasts | CALD1, CYR61, TAGLN, MT1X, PRELP, TPM2, GPX3, CTGF, IGFBP5, SPARCL1 |
| B cells | CD69, CORO1A, LIMD2, BANK1, LAPTM5, CXCR4, LTB, CD79A, CD37, MS4A1 |
| Mast cells | VWA5A, RGS13, C1orf186, HPGDS, CPA3, GATA2, MS4A2, KIT, TPSAB1, TPSB2 |
| Interstitial Mφ perivascular | MRC1, RNASE1, FGL2, RNASE6, HLA-DPA1, GPR183, CD14, HLA-DPB1, MS4A6A, AIF1 |
| SMG mucous | FKBP11, TCN1, GOLM1, TFF3, PIGR, KLK11, MARCKSL1, CRACR2B, SELM, MSMB |
| AT2 proliferating | CDK1, LSM3, CKS1B, EIF1AX, UBE2C, MRPL14, PRC1, CENPW, EMP2, DHFR |
| Goblet (subsegmental) | MDK, MUC5B, SCGB1A1, CP, C3, TSPAN8, TFF3, MSMB, PIGR, BPIFB1 |
| Pericytes | MYL9, SPARC, SPARCL1, IGFBP7, COL4A1, GPX3, PDGFRB, CALD1, COX4I2, TPM2 |
| SMG duct | PIP, ZG16B, PIGR, SAA1, MARCKSL1, ALDH1A3, SELM, LTF, RARRES1, AZGP1 |
| Mesothelium | CEBPD, LINC01133, MRPL33, UPK3B, CFB, SEPP1, EID1, HP, CUX1, MRPS21 |
| SMG serous (nasal) | ZG16B, MUC7, C6orf58, PRB3, LTF, LYZ, PRR4, AZGP1, PIGR, RP11-1143G9.4 |
| Ionocyte | FOXI1, ATP6V1A, GOLM1, TMEM61, SEC11C, SCNN1B, ASCL3, CLCNKB, HEPACAM2, CD24 |
| Alveolar Mφ MT-positive | GSTO1, LGALS1, CTSZ, MT2A, APOC1, CTSL, UPP1, CCL18, FABP4, MT1X |
| Fibromyocytes | NEXN, ACTG2, LMOD1, IGFBP7, PPP1R14A, DES, FLNA, TPM2, PLN, SELM |
| Deuterosomal | RSPH9, PIFO, RUVBL2, C11orf88, FAM183A, MORN2, SAXO2, CFAP126, FAM229B, C5orf49 |
| Tuft | MUC20, KHDRBS1, ZNF428, BIK, CRYM, LRMP, HES6, KIT, AZGP1, RASSF6 |
| Plasmacytoid DCs | IL3RA, TCF4, LTB, GZMB, JCHAIN, ITM2C, IRF8, PLD4, IRF7, C12orf75 |
| T cells proliferating | TRAC, HMGN2, IL32, CORO1A, ARHGDIB, STMN1, RAC2, IL2RG, HMGB2, CD3D |
| Subpleural fibroblasts | SERPING1, C1R, COL1A2, NNMT, COL3A1, MT1E, MT1X, PLA2G2A, SELM, MT1M |

| | |
|---|---|
| Lymphatic EC proliferating | S100A16, TUBB, HMGN2, COX20, LSM2, HMGN1, ARPC1A, ECSCR, EID1, MARCKS |
| Migratory DCs | IL2RG, HLA-DRB5, TMEM176A, BIRC3, TYMP, CCL22, SYNGR2, CD83, LSP1, HLA-DQA1 |

### SUPPLEMENTAL DATABASES

**Supplemental Database S1: Spatialomics Gene Expression data for LAM_D1 supporting Figures 2-3.** A database containing significantly upregulated genes per cluster for LAM-lung tissue donor LAM_D1 spatialomics.

**Supplemental Database S2: Spatialomics Gene Expression data for LAM_D2 supporting Figures 2-3.** A database containing significantly upregulated genes per cluster for LAM-lung tissue donor LAM_D2 spatialomics.

**Supplemental Database S3: Spatialomics Gene Expression data for the Combined LAM lung Tissues supporting Figures 2-3.** A database containing significantly upregulated genes per cluster for the combined dataset of both LAM-lung tissue donor spatialomics.

**Supplemental Database S4: Phospho Kinase Array supporting Figure 5A.** Raw data and analyzed expression for the phosphor kinase array performed on LAM-cells in the presence of vehicle or 7µM Sorafenib.

**Supplemental Database S5: Spatialomics IPA Canonical Pathway Enrichment for LAM-Core.** A database containing the significant differentially expressed pathways analyzed through IPA for the genes in the LAM-Core cluster.

**SUPPLEMENTAL FIGURES**

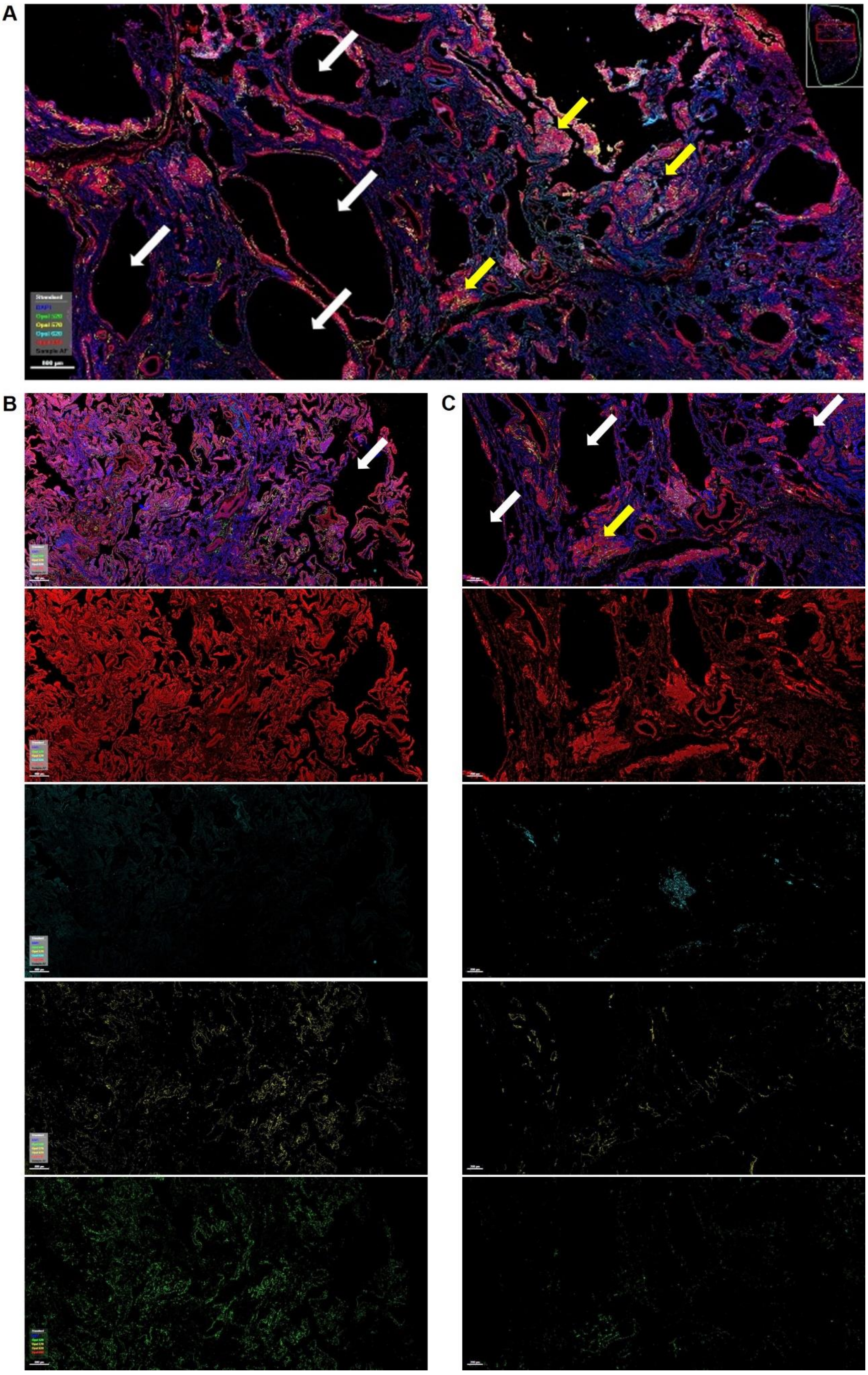


**Supplemental Figure S1: Identification of LAM nodules in patient LAM lung tissue**. Representative images of LAM lung tissues stained with VEGFR3 (green), PDPN (yellow), HMB-45 (cyan) and SMA (red). Nuclei are counterstained with DAPI (blue) in all images. White arrows indicate alveolar cysts and yellow arrows highlight LAM nodules forming near cysts. Scale bars in all images are 800 µm.

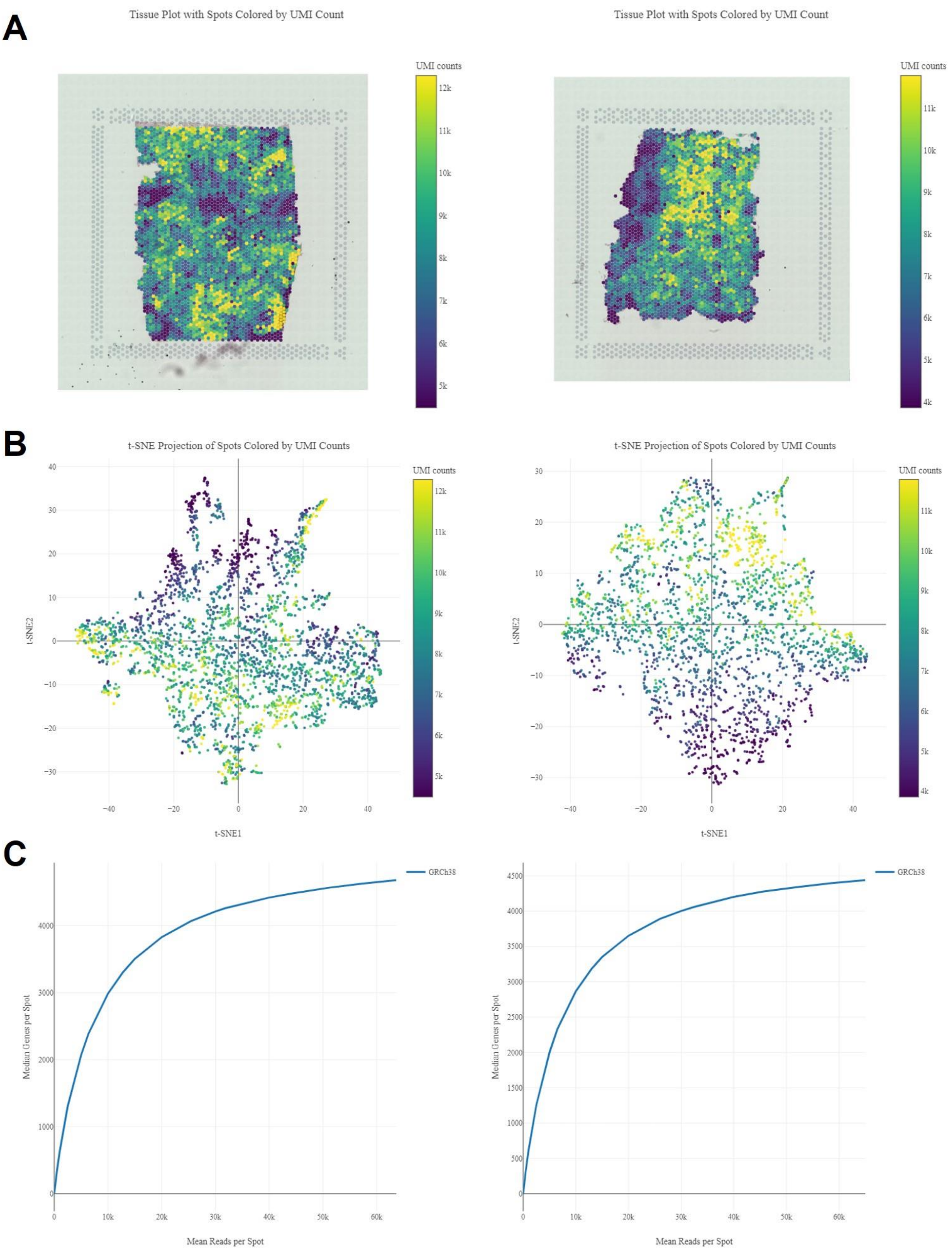


**Supplemental Figure S2: Quality Control analysis of Spatial-Transcriptomics Sequencing.** Tissue plots for LAM_D and LAM_E 2 lung sections showing UMI counts per spot mapped to the tissue (A) and to the tSNE plots (B). C) Mean genes per spot mapped to mean reads per spot for LAM_D1 and LAM_D2.

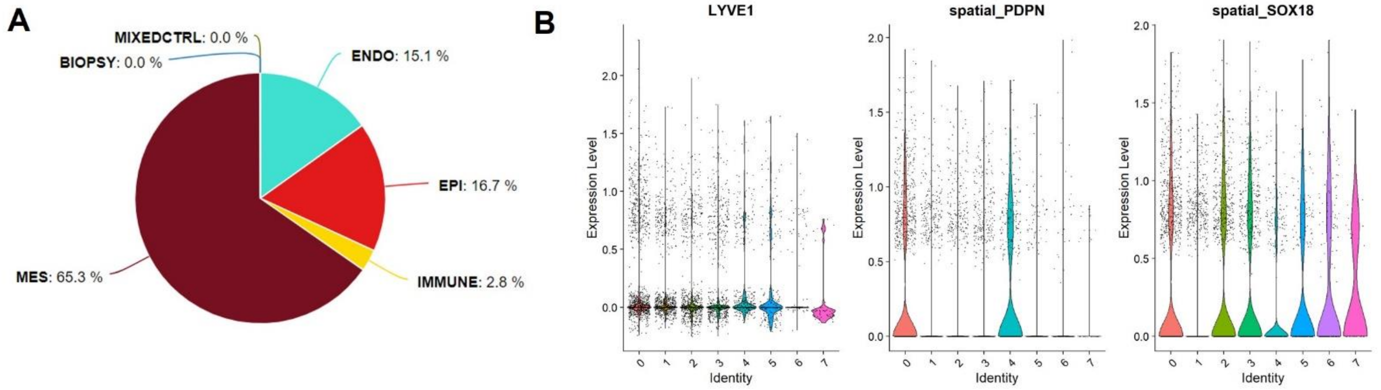


**Supplemental Figure S3: Contribution of LEC to the LAM-Core cluster.** (A) Pie chart of the mapped cell types reflecting the gene expression in the cluster representing the LAM core (Cluster 4). (B) Gene expression profile visualized as violin plots for each cell cluster for core LEC genes LYVE1, PDPN, SOX18 for combined datasets from LAM_D1 and LAM_D2. LAM Core cluster is identified as cluster 4 (turquoise).

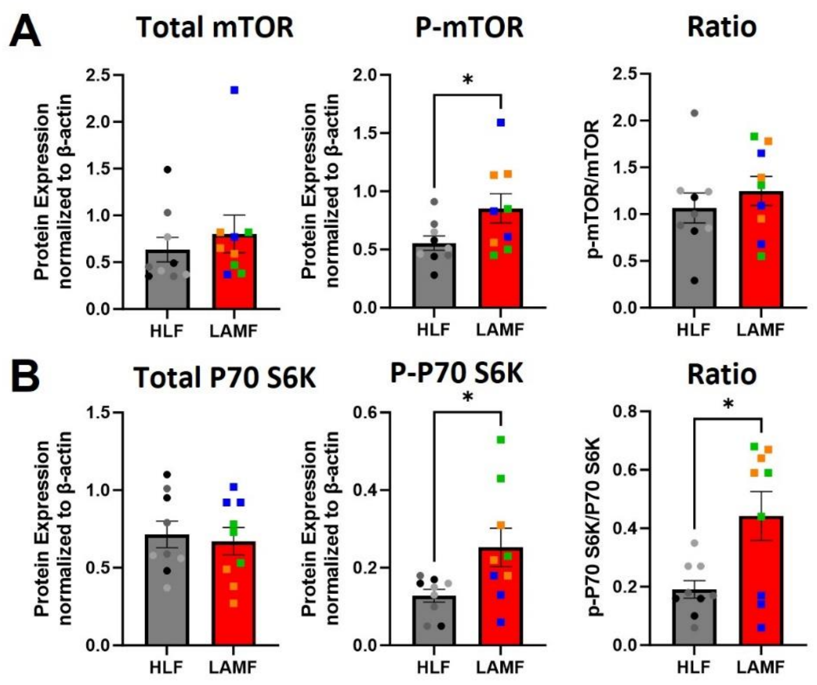


**Supplemental Figure S4: Characterization of HLF and LAMF.** A-B) Protein expression for Total and phosphorylated mTOR (A) and p70-S6K (B) and the phosphor/total protein expression ratios for HLF (grey) compared to LAMF (red). The coloured dots represent different donors each with 3 experimental repeats (n=9. Data shown represents mean±SEM, significance is represented by *p<0.05, N=3 independent donors per cell type.

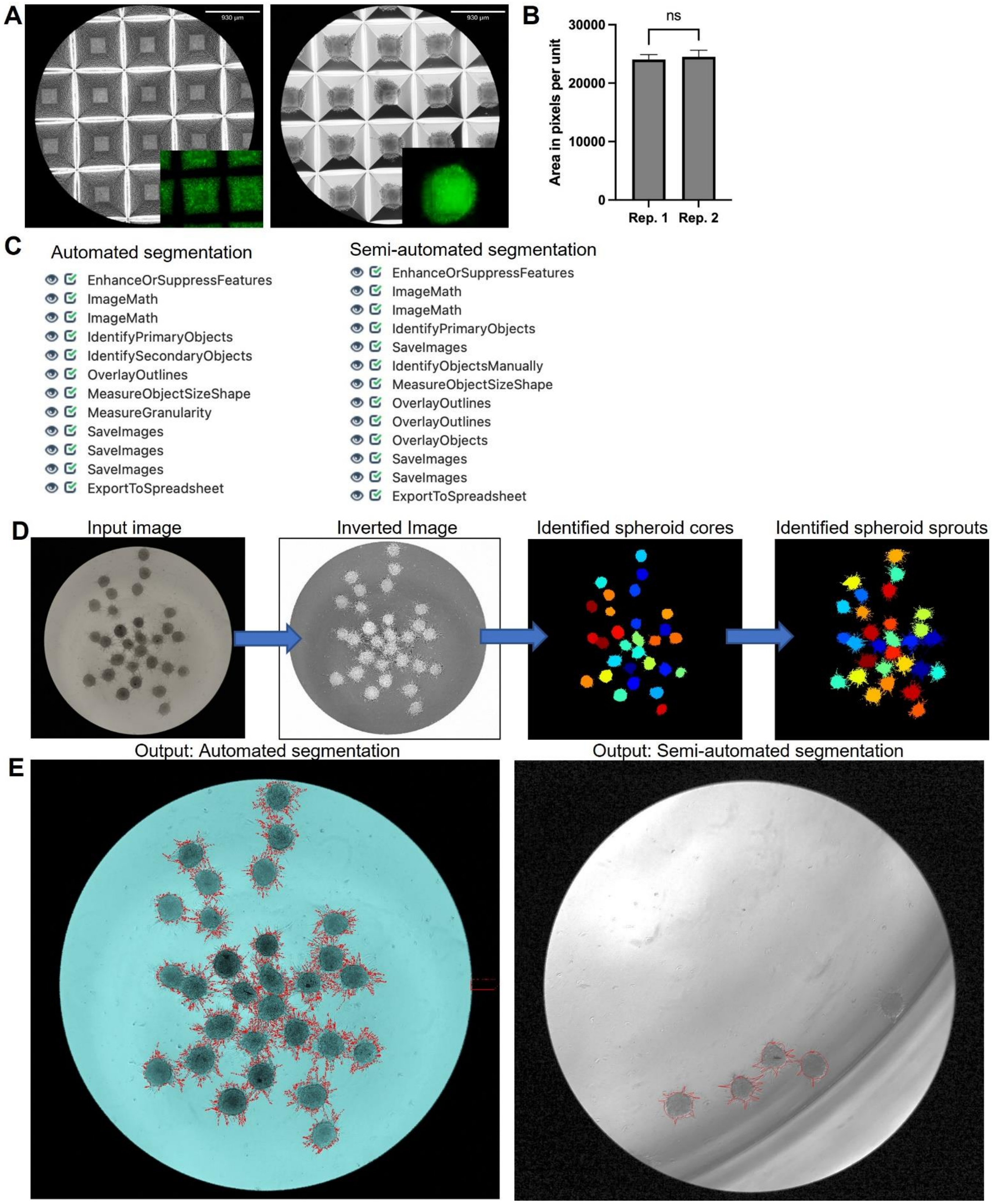


**Supplemental Figure S5: Cell Profiler command pipeline and Image segmentation.** (A) Representative images of CellTracker-Gren Cells before (time 0hr) and after spheroid formation (24hrs) in Aggrewell plates. (B) Comparison of spheroid size for two replicate experiments using LAMF. (C) Analysis command pipeline used in this study for automated versus semi-automated image segmentation. (D) Representative images of the pipeline steps starting with an inversion of the image with enhancement of line structures, in this case sprouting. Image masks automatically generated by cell profiler segmenting spheroid cores and spheroid sprouts. (E) Outlines of spheroids detected by the automated pipeline versus semi-automated pipeline.

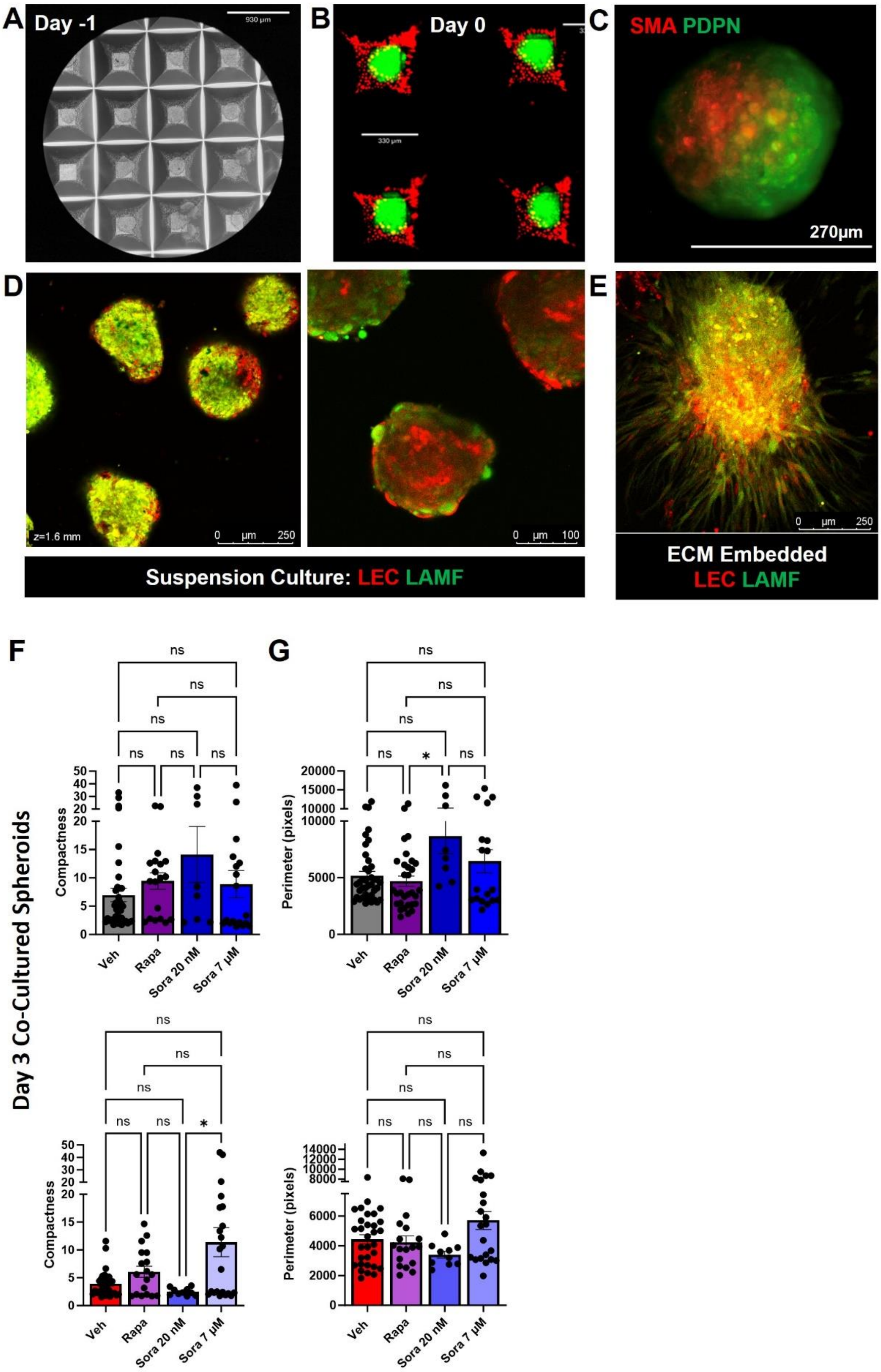


**Supplemental Figure S6: Co-Cultures LAMF-LEC spheroids and early 3-D invasion into the extracellular matrix.** Representative images of cells plated in aggrewell plates (A), addition of CellTracker-Red LEC to CellTracker-Green LAMF (B), Day 1 co-cultured spheroids stained with SMαA (LAMF) and PDPN (LEC) (C), suspended co-cultured spheroids after 24 hours labelled with Cell-Tracker (D) and CellTracker labelled co-cultured spheroids after 7 days in ECM culture (E). (F-G) Relative comparison of compactness (F) and Perimeter (G) for co-cultures spheroids after 3 days of 3-D culture in ECM. Data shows vehicle (grey/red) compared to Rapa (purple) and 20nM Sora (dark blue) and 7µM Sora (light Blue) after 3 days for HLF (top) and LAMF (bottom). Each dot represents a single spheroid. Data shown represents mean±SEM, significance is represented by *$p<0.05$, N=3 independent donors per cell type.
